# CoSpred: Machine learning workflow to predict tandem mass spectrum in proteomics

**DOI:** 10.1101/2024.01.18.576149

**Authors:** Liang Xue, Shivani Tiwary, Mykola Bordyuh, Pierre M Jean-Beltran, Robert Stanton

## Abstract

In mass spectrometry-based proteomics, the identification and quantification of peptides and proteins is usually done using database search algorithms or spectral library matching. The use of deep learning algorithms can help improve the identification rates of peptides and proteins through the generation of high-fidelity theoretical spectrum which can be used as the basis of a more complete spectral library than those presently available. Current methods focus on predicting only backbone ions, such as y- and b-ions. However, the inclusion of non-backbone ions is necessary to truly improve spectral library matching. Here we focus on providing a user-friendly machine learning workflow, which we call **Co**mplete **S**pectrum **Pred**ictor (CoSpred). Using CoSpred users can create their own machine learning compatible training dataset and then train a Machine Learning model to predict both backbone and non-backbone ions. For the model a transformer encoder architecture is used to predict the complete MS/MS spectrum from a given peptide sequence. This model does not require background knowledge of fragment ion annotations or fragmentation rules. The model outputs the set of pairs (*M*_*i*_, *I*_*i*_) where *M*_*i*_ is the m/z (mass-to-charge ratio) of a peak in the spectrum and *I*_*i*_ is the intensity of the peak. The model presented here for validation was trained on the dataset available in the MassIVE data repository and shows superior performance in terms of various metrics (e.g. precision/recall for mass, cosine similarity for peak intensity, etc) between the true and predicted spectra. Furthermore, CoSpred can be used to create custom models that allow for accurate spectrum prediction for different experimental conditions. In addition to the transformer model provided in the package, the code is built modularly to allow for alternate ML models to be easily “plugged in”. The CoSpred workflow (preprocessing->training->inference) provides a path for state-of-art ML capabilities to be more accessible to proteomics scientists.

## INTRODUCTION

### Mass Spectrometry Spectrum Prediction

Shotgun proteomics^1, 2^ is a widely used method to identify and quantify proteins in samples of interest (e.g., cells and tissues). Digested peptides are fragmented in a mass spectrometer producing a fragment spectrum (MSMS). The fragment spectra show some regularity^3^ due to the oligomeric structure of peptides and the predictability of peptide bond breakage patterns. This can be exploited to interpret the spectra and determine the sequence of amino acids as well as any covalent modifications of the amino acids. Knowing the physical method of fragmentation (for instance collision-induced dissociation (CID)^4, 5^, higher-energy collisional dissociation (HCD)^6^ or electron transfer dissociation(ETD)^7^) the masses of the dominant peptide fragments can be calculated from the sequence. Classic peptide database search engines use the theoretical set of fragment masses to identify the peptides in the sample^8-10^.

Spectral library matching is another widely used approach for peptide identification. This method incorporates the fragment intensity information, in addition to the discrete set of expected fragment masses. Intensity information is taken from previously measured spectral libraries and improves sensitivity and specificity of peptide identification^11-13^. As the method relies on the spectra of known sequences it is not able to identify novel peptides for which fragment spectra have not been acquired. The creation of spectral libraries can be sample specific, expensive and time consuming, therefore in-silico generated spectra using machine learning methods are an attractive alternative^14-16^.

Over the last two decades many classical machine learning methods have been used to predict peptide fragment spectra,^17, 18^ and more recently deep learning methods^16, 19, 20^ have been developed which can predict peptide fragmentation spectra from the amino acid sequence with the accuracy of experimental replicates^16, 19^. The most popular deep learning methods such as Prosit^19^, DeepMass:Prism^16^ and pDeep^20^, use recurrent neural networks (RNN)^21^ to model the fragment spectrum of y- and b-ion series. However, different fragmentation protocols lead to the generation of distinct *a/b/c/x/y/z* ion series, which still only account for about 70% of the experimental high intensity peaks. The remaining peaks which come from non-backbone ions are not matched by current peptide search algorithms, leaving room for improvement in peptide identification accuracy and precision ^22^. Here, the a/b/c/x/y/z ion series denote backbone ions and remaining peaks non-backbone ions.

Complete spectrum prediction methods using deep learning architectures make predictions independent of fragmentation rules and annotations. PredFull^22^ predicts complete spectrum, both backbone and non-backbone ions using convolutional neural networks (CNN). for higher energy collisional dissociation (HCD) spectra, it uses two CNN models for precursor charge 2+ and 3+ separately. To include precursor charges 1 and 4 they use multi-task learning. Here, we aim to create a single model for all precursor ion charges using a transformer architecture. Recently, transformer architectures have been successfully applied to natural language processing tasks. They have the potential to identify long range dependencies in the sequence and learn relationships between the amino acids in a sequence irrespective of their positions. Transformer architectures use multi-head attention layers to learn patterns and all the calculations are done in parallel. Peptide spectrum prediction has been shown to be affected by long range interactions as the propensity of peptide bond cleavage can be determined by a combination of the local molecular environment and more distant sequence effects.^23^ Compared to RNN/CNN which are limited in the modelling capacity due to the way how the data is internally processed sequentially, transformer architectures are now gaining interests of being used for proteomics. Previous studies like Casanovo have used a Transformer encoder/decoder architecture similar to this application where the input is a peptide sequence and output the next amino acid in the sequence.^24^ Such a workflow was used for prediction of a peptide sequence from a given MSMS spectrum.

Here, we developed a workflow named **Co**mplete **S**pectrum **Pred**ictor or **CoSpred**, where a user can convert raw files, independent of mass spectrometry instrument vendor, and convert them into machine readable tensors, which later are used as input to train transformer encoders for complete MSMS spectrum prediction. The complete MSMS predictor is independent of fragment ion annotations and their fragmentation rules, and instead predicts the (*M*_*i*_, *I*_*i*_) pairs, where *M*_*i*_ is the m/z (mass-to-charge ratio) of a peak in the spectrum and *I*_*i*_ is the intensity of the peak^23^. CoSpred creates a user-friendly workflow for data preprocessing from raw file formats found in proteomics data repositories such as PRIDE^25^ and MassIVE-KB^26^. These raw files are then converted into mzML and mgf file formats. Combining peptide identification results from software like MaxQuant^27^ and Proteome Discover (Thermo Scientific) with mzML/mgf files a text compatible file like HDF5 is created containing m/z and intensity values for each peptide. This dataset is then divided into training, validation and test sets which are used for model training. The best model is saved and used for complete spectrum prediction of the test set. Prediction results can be saved into different output formats such as mgf, msp, csv and a long format suitable for creating an in-silco spectral library that can be used in various applications. The most common expected use is for the predicted spectral library in data independent acquisition (DIA) analysis software such as MaxDIA^14^, DIANN^15^ and Spectronaut (Bruker Biognosis). This workflow is designed for compatibility with multiple possible ML architectures. In this paper, we demonstrate how to plug in existing mature architectures like Prosit, and show an end-to-end workflow example starting from raw mass spectrometry files, followed by training/fine-tuning, spectrum prediction, and finally loading a predicted library into DIANN for classical DIA data processing^15^.

## METHODS

### CoSpred workflow

The entire CoSpred workflow is divided into three sections, first data preprocessing of the mass spectrometry datasets, second a transformer model created using the PyTorch framework and lastly the prediction of spectra using saved models. The workflow is shown in **Figure 1** and described in the following sections.

**Figure 1.**
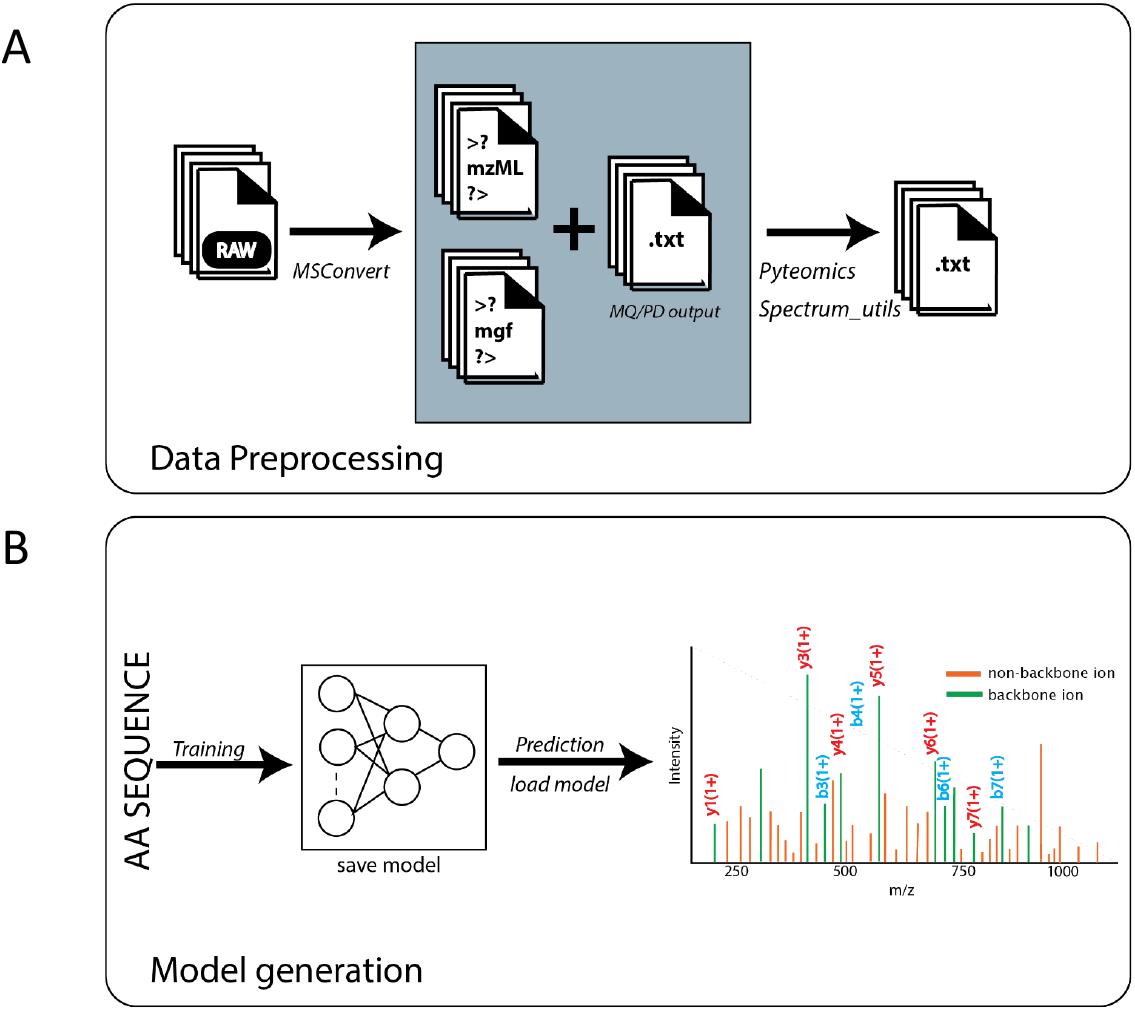
CoSpred workflow. A) Shows the workflow of data preprocessing. To make data preprocessing user friendly and accessible we provide steps to convert raw files into xml formats such as mgf and mzML and by combining this with the output of MaxQuant^27^ or Proteome Discoverer the identified peptides, m/z ratio and intensities are all saved into a csv file, which is later used as the input for any machine learning model. B) Amino acid sequence and precursor charge are taken from the input files and converted into tensors which are used as input for machine learning models, the model with least validation loss is selected and saved for future use. Predictions can then be made of complete spectrum including non-backbone as well as backbone ions.

### Dataset Preprocessing

In the data preprocessing step, we start with acquiring raw files from mass spectrometry instruments that have been deposited in a proteomics database repository like PRIDE^25^ and/or MassIVE^28^. These raw files are instrument specific and require conversion to a common format. MSConvert ^29^ is used to convert these raw files into open formats such as mzML and mgf. These file formats conform to Proteomics standards and easily readable. The raw files do not contain identified peptides and must be paired with the output text from peptide identification software such as MaxQuant^27^ or Proteome Discoverer (Thermo Scientific). The identified peptide text file or xml files are parsed to get the m/z ratio and its corresponding intensity peak. Pyteomics^30^ and Spectrum_utils^31^ were used to parse the masses and corresponding intensity peaks and save to an annotated text file like Mascot Generic Format (MGF)^32^ format as shown in **Figure 1A**. The full data set is then divided into training, validation, and test sets, which are later used as input for the machine learning module.

As an example, we used the raw files and Proteome Discoverer (Thermo Scientific) output files (PSMs.txt) from ProteomeTools^33^ through the PRIDE^25^ database. Raw files were converted into both mzML and MGF formats using MSConvert^29^. For each Peptide spectrum match (PSM), all the masses and intensities were parsed from the MGF files, along with the scan number, sequence, modified sequence, precursor charge, retention time, and collision energy using the Pyteomics^30^ package in python and then saved in an hdf5 file. This file was used as the input for the training module, where sequence, precursor charge and collision energy and intensity peaks^23^ were converted into torch tensors to be used as input in the PyTorch framework. Alternatively, users could also opt to use a Tensorflow framework considering some published models were developed through Tensorflow, such as Prosit or pDeep. The detailed description of how to run the data preprocessing workflow to create a training set from experimental data is given in the readme file in the GitHub repository (https://github.com/pfizer-opensource/CoSpred). One crucial step included in the workflow is the filtering of input data to get unique peptide spectrum matches (PSMs) for each precursor charge and collision energy from the raw files. It is also important that the random division of data into training, test and validation results in equal charge and sequence length distributions among the dataset splits. To get the annotations of y-, b-ions up to 3+ ion charges we used the Spectrum_utils^31^ package. This is also used to compare CoSpred results of backbone ions y-, b-ion with the existing spectrum predictors^16, 19, 20^ results. Peptide sequence is converted into a sequence integer and the maximum length of sequence was fixed to be less than 30. Charge was taken as an integer from 1 to 6 and collision energy was normalized to be from 0 to 1 giving a float value.

### Transformer architecture for complete MSMS spectrum prediction

A transformer encoder architecture was used to predict the complete MSMS spectrum. The model consists of a transformer encoder as shown in **Figure 2**. The encoder takes the peptide sequence as input and outputs a one-dimensional vector consisting of pairs *M*_*i*_,*I*_*i*_, where *M*_*i*_ is the *m/z* (mass-to-charge ratio) of a peak in the spectrum and *I*_*i*_ is the measured intensity of the peak. The 1-dimensional vector generated is of length 3000 with a bin width size of 0.5 Dalton, resulting in a maximum m/z ratio of 1500. The maximum m/z can be changed by the user if the dataset contains spectra greater than 1500 m/z. The bin size width can also be modified according to the needs of the user. Metadata such as charge, and collision energy are included as a separate embedding layer. Input and metadata embedding layers were then concatenated and sent to a transformer encoder block within PyTorch. The positional encoding was fixed as sin and cosine functions.

**Figure 2:**
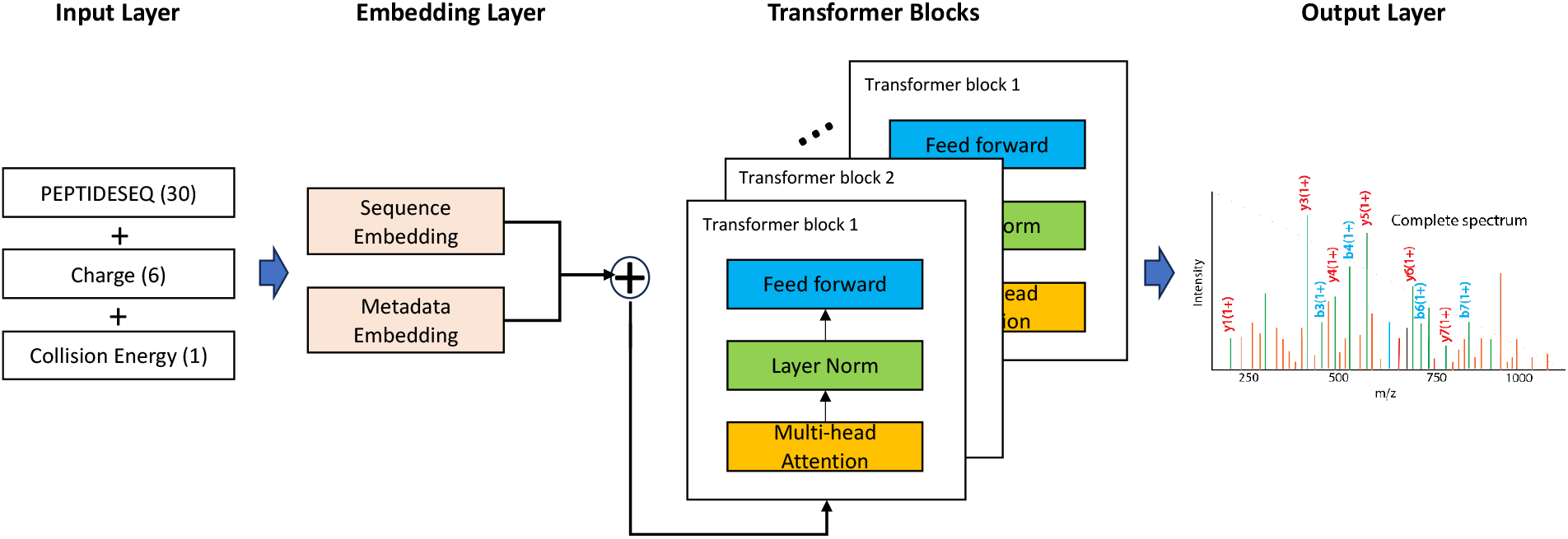
Transformer Architecture. The input of this architecture is peptide sequence represented as sequence integer and charge (integer) and normalized collision energy (float). Sequence and metadata (charge and collision energy) are embedded separately and then concatenated together. The embedding layer was given as input to the Transformer encoder layer, which consisted of 8 layers and 16 attention heads, and the feed forward network gave output as a 1 dimensional vector of length 3000.

### Training Hyperparameters

We trained a model with an embedding size of 256, 8-layers, and 16 attention heads. A batch size of 1024 PSMs was selected with a weight decay of 0.1. A learning rate of 0.001 was taken. The model was trained until convergence. Spectral contrast loss^34^ was used to identify the best model. Other metrics such as cosine similarity, precision, recall, F1 score, etc were also used to evaluate the training performance. The model with lowest validation loss was saved and used for testing. The spectral angle loss was calculated using following equation:

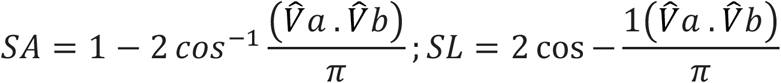

where

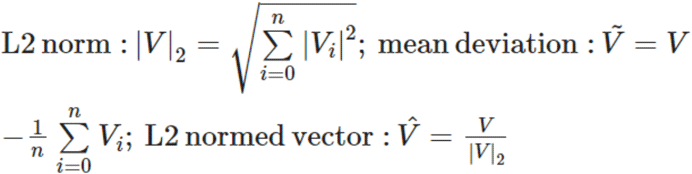

### Inference

The saved model can be used to predict complete spectrum for new peptide sequences. The predicted output can be written into various formats such as mgf, plain csv or tab separated files. We also created DIANN^15^ long output format with all the required columns. These files then can be used directly in generating *in silico* spectra libraries for DIA data analysis for example in the DIANN^15^ or MaxDIA^14^ software. Detailed steps are provided in the ReadMe page on the CoSpred GitHub repository.

### Datasets

Training datasets (**Table 1**) consist of the ProteomeTools synthetic peptides^33^ with both b/y ion annotated HDF5 format as well as raw MGF format from the MassIVE^26^ repository. In addition, the extended synthetic peptides library (HCD spectra generated in the ProteomeTools and Bioplex experiments) was also retrieved from MassIVE-KB spectral library v2 full release. The spectra were filtered with precursor charges ranging from 1 to 6. A study of the effect of dataset size on model training is shown in **Figure 3**. The charge and length distribution of the training dataset is shown in **Supplementary Figure S1**. The data was randomly indexed and divided into training, validation, and test sets at a ratio of 90:5:5.

**Table 1:** Summary of datasets used for the study. High quality HCD spectra for synthetic peptides were retrieved from three resources, which were the original ProteomeTools HCD Spectral Library in MGF format, the Prosit provided HDF5 dataset, and the combined ProteomeTool and Bioplex HCD spectra library in MGF format retrieved from MassIVE. Training and validation sets were used for training the architecture to tune the hyperparameters and the test set was used later to evaluate the models. The “Number of Spectra” column indicates the number of spectrum records that were used in this project after preprocessing and filtering, while in parenthesis is the number of records in the whole dataset according to the original MassIVE resources.

| Source | Library Name | Format | File Size | Number of Spectra | Precursors | Peptides |
| --- | --- | --- | --- | --- | --- | --- |
| Gessulat et.al.<sup>35</sup> | Prosit datasets | HDF5, b-/y-ion annotated | 5.23GB | 6,787,933 (21,764,501) | 576,256 | 549,375 |
| MassIVE | ProteomeTools HCD Spectral Library | MGF | 1.1GB | 262,930 | 264,836 | 204,625 |
| MassIVE | KB 2.0.15, synthetic | MGF | 6.23GB | 1,062,010 | 1,662,275 | 889,204 |

**Figure 3:**
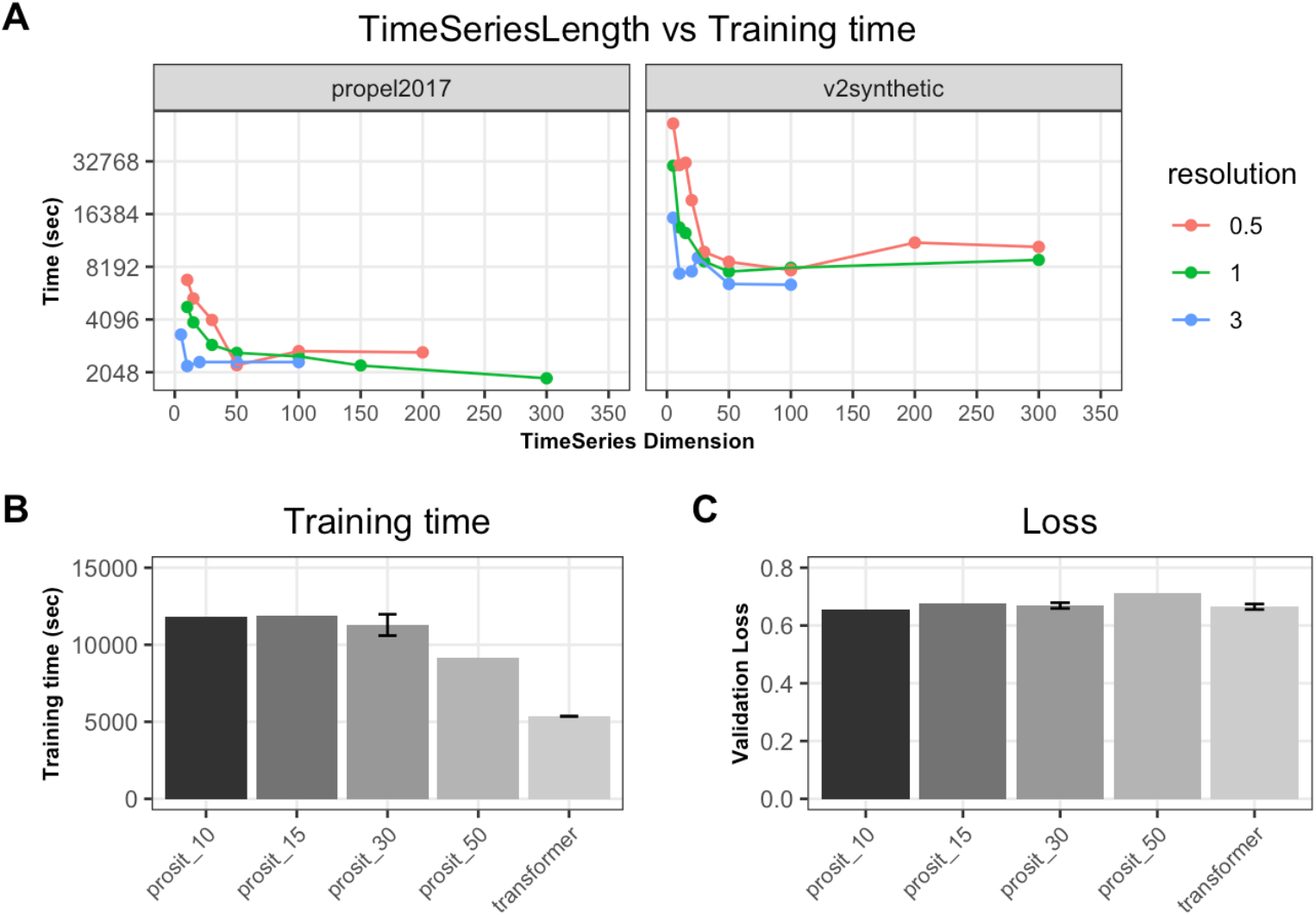
Training time and loss for various Prosit settings. A) Comparison of different time step numbers in the last layer of the Prosit model with respect to various mass resolutions and datasets. B) Training time for the different Prosit architectures given a fixed training set with 200k spectra. C) Same layout but with the loss measurement.

## RESULTS

### Speed up of full spectrum prediction model through CoSpred workflow

While Bidirectional Gated Recurrent Unit (BiGRU)^36^ recurrent neural network models like Prosit have been proven to successfully predict MS spectrum at high fidelity^35^, it is not clear if such architectures still perform optimally for full spectrum prediction. To test this hypothesis, we first compared a handful of hyper-parameter combinations in terms of their converged loss and training speed.

Independent of datasets, we observed that an increase of dimensions on the time series layer of the Prosit architecture improves the training speed significantly (by up to 50x) (**Figure 3A**). Interestingly, beyond that elbow point, further increasing the dimensions didn’t improve training speed. Also, examining various spectrum resolutions including 3Da, 1Da, 0.5Da showed correlation between higher prediction resolution and longer training time. However, such speed differences diminished when time steps increased above 50. Given these results and for practicality, all our following experiments were done using a 0.5Da mass resolution.

Next, we compared the training time as well as converged loss between the transformer architecture and various Prosit settings. To better balance between training time and loss, we chose 30 dimensions for the last time step layer of the Prosit model, which is shown as Prosit_30 in **Figures 3B** and **3C**. Consistently, the transformer based CoSpred model yields faster training given the same number of epochs (**Figure 3B**), while the final loss measured by spectral distance is comparable among all models (**Figure 3C**). The results suggest that transformer architectures may be a viable alternative ML strategy for full MS spectrum prediction alongside methods like Prosit.

### Training transformer model with various size of spectra datasets

To test the accuracy of the model depending on the size of the dataset, the transformer encoder was trained for datasets ranging from 10,000 to around 7 million spectra. For validation we used the spectral distance as the loss between true and predicted spectrum, shown in **Figure 4**. As shown in this figure, there was improvement in spectral distance with increased data set size. For a simpler task, predicting the b/y ion fragmentation spectra, the loss significantly decreased when the training set size was increased from 10,000 to 1 million, at which point it appeared to reach a plateau (**Figure 4A**). On the other hand, for the full spectra prediction, the loss for both the CoSpred and Prosit were still monotonically decreasing up to the full MassIVE KB 2.0.15 synthetic dataset (v2synthetic) which contains around 1 million spectra (**Figure 4B**), indicating there is still potential to further improve the model with larger sets of data.

**Figure 4:**
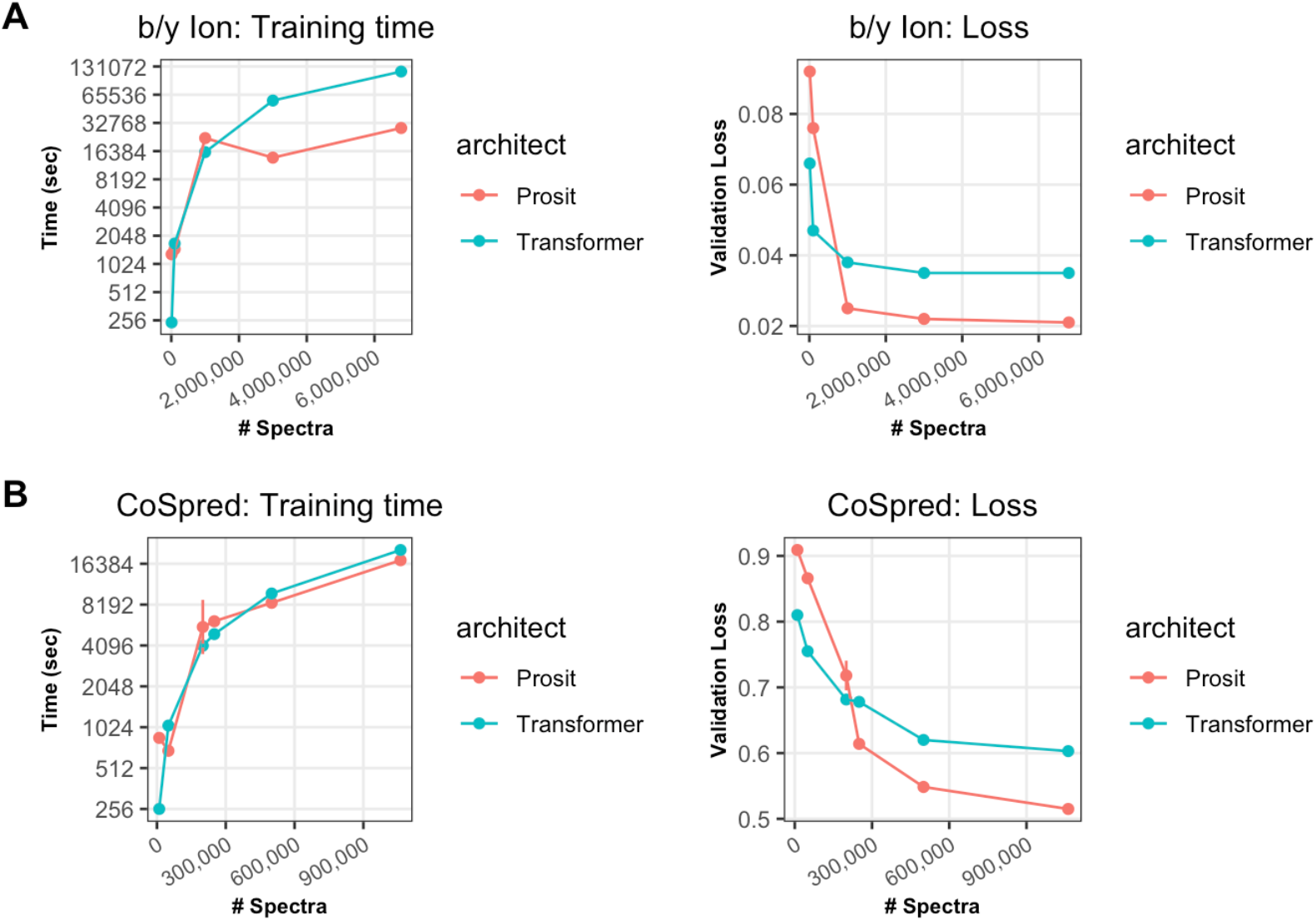
Training Performance for the CoSpred Transformer based architecture. A) Comparison of dataset size. The plot shows training datasets of different sizes ranging from 10K to 7 million on the x axis, and on the y axis is the training time until convergence (left panel), as well as the spectral distance loss between true and predicted b/y ion spectra (right panel). B) Same layout as with the prediction being for the full spectrum instead of b/y ions.

When investigating the performance of the model, we didn’t observe significant bias towards peptide charge states (**Supplementary Figure S2A)**. On the other hand, it appears that the accuracy of prediction decreases when the peptide length increases (**Supplementary Figure S2B**). We suspect that this might be due to sparse training examples for longer peptides, especially as string diversity increases with the longer sequence length.

### Hyper-parameter tuning of spectrum prediction model through the CoSpred workflow

With a better understanding on the baseline performance of both the RNN and Transformer based ML architectures on full spectrum prediction, we further executed hyperparameter tuning by grid searching on the number of layers, number of heads, and how an early stop (i.e. restricting the maximum epoch number) influences the model performance. For the grid search, 200k spectra from v2synthetic dataset with 4:1 split between training and validation was used. The final input training dataset was pre-filtered with precursor charges ranging from 1 to 6. 1000 spectra were held-out for testing.

Compared to a baseline Prosit model, the CoSpred model with a maximum epoch number of 100 showed superior performance with about 2x faster training time and lower validation loss. Further increasing the complexity of the transformer model, such as through the inclusion of more layers, increased numbers of attentions heads, or adding a convolutional layer in addition to multilayer perception (MLP), like published in the GPT2 model^37^, all dramatically increased the training time since the ultra-high number parameters slows the model convergence. In addition, lifting the early stop criteria and letting the model strictly converge also doubles the training time. Most of the slower methods gave marginal benefit for validation loss improvement. As a result, we suggest for this task within CoSpred 8 layers and 16 heads to be an optimal model to move forward.

### Transformer based architecture improves fragment mass and intensity prediction

With the choice of hyper-parameters (full list presented in **Supplemental Table S1**), the Prosit and CoSpred models were trained with the Prosit dataset provided by Gessulat et.al.^35^ for b/y ion prediction, as well as the “v2synthetic” dataset for full spectrum prediction. The best model was saved where the validation loss had converged to a minimum. Using a 1000 spectra holdout set from either dataset for testing purposes, the precision-recall curve of the model for the training data is shown in **Supplemental Figure S3**. A full summary of the performance metrics is shown in **Table 2**.

**Table 2:**
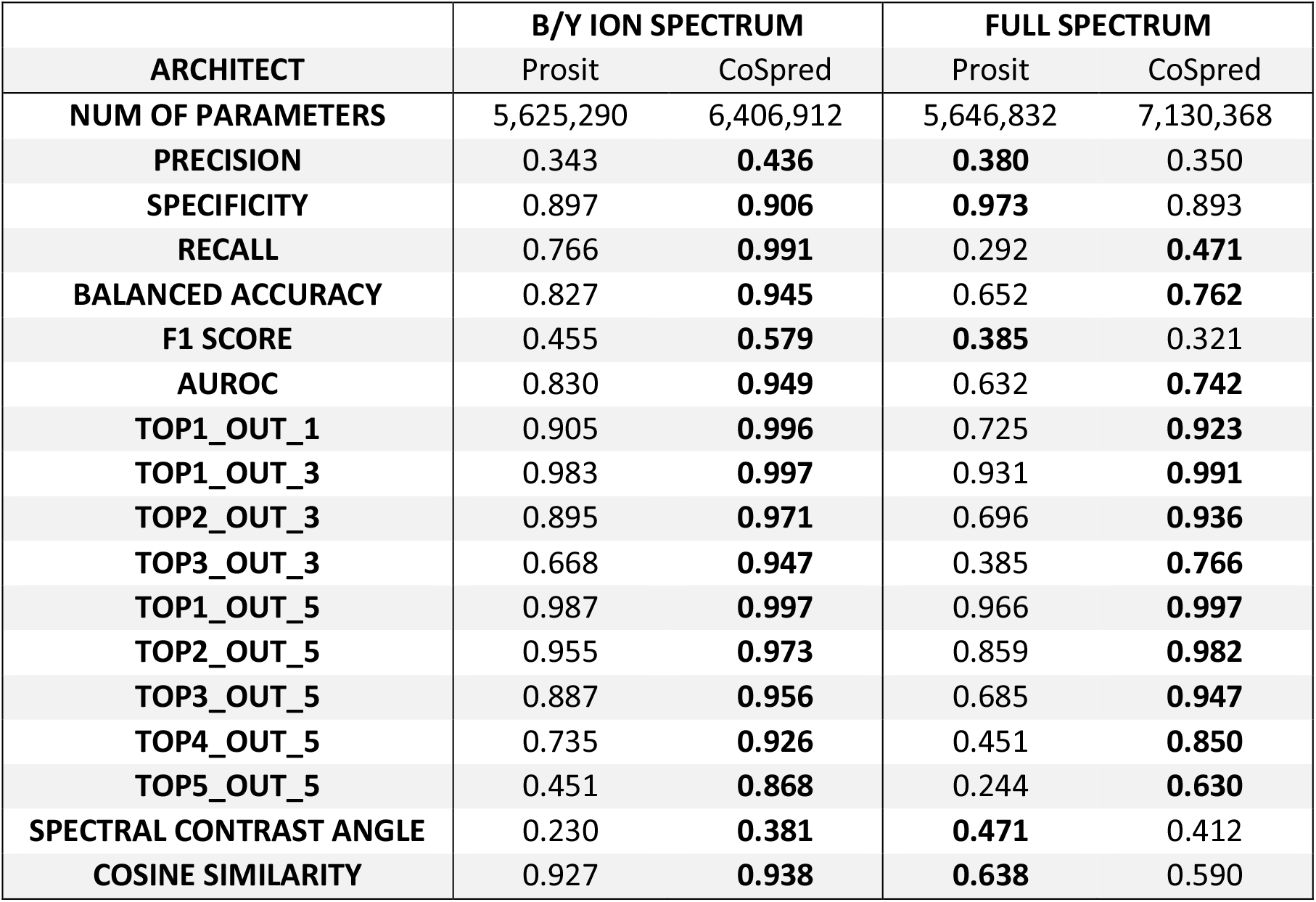
Metrics for testing performance of CoSpred vs Prosit in two prediction tasks. Bold indicates the superior methods between the two architectures for each measurement per application. Note: Precision = (True Positive)/(True Positive + False Positive). Recall = (True Positive)/(True Positive + False Negative). Specificity = (True Negative)/(True Negative + False Positive). Balanced Accuracy = (Recall + Specificity)/2. F1 score is the harmonic mean of Precision and Recall. AUROC is Area under curve of Receiver Operating Characteristic (ROC) curve. TopX_OUT_Y is the probability of predicted top X intensive peaks were also the top Y peaks in ground truth. Spectral Contrast Angle was originally proposed by Wan et.al.^34^ Cosine Similarity was computed by CosineSimilarity function in Pytorch package.

There are two major categories of performance measurements shown in the table. Spectral Contrast Angle and Cosine Similarity consider the peak intensity similarity between reference and predicted spectrum, while the rest were focused on binary prediction of whether certain peaks existed in the expected m/z according to the reference ground truth. In essence, when considering the whole mass spectrum as 3000 mass bins with 0.5Da window size, we framed the full spectrum prediction as a multiclass classification task, so that existing ML metrics like precision and recall could be applied for evaluation.

For an application like b/y ion prediction, the transformer based CoSpred model appeared to outperform on all metrics, including both mass and intensity prediction. Some example spectra are visualized in **Supplemental Figure S4**. In the case of a more complex task like full spectrum prediction, we observed a tradeoff between precision and recall for the two architectures. Prosit was more conservative with higher precision in generating peaks, while CoSpred could recover more peaks in prediction, leading to higher recall / sensitivity but lower precision. Some representative cases are shown in **Supplemental Figure S5**. The highly imbalanced nature of such spectrum data with 3000 classes, is due to the fact that most mass windows have no signal / peak, we suspect some performance indicators like the F1 score would be biased against the less precise CoSpred model compared to less sensitive Prosit model. Moreover, we anticipated that higher sensitivity may be more beneficial practically, for cases such as setting up target mass spectrometry methods when the researcher needs to monitor top intensity fragmentation ions. In such a case, we observed the top 5 intensity peaks with a correct ranking was better preserved in the CoSpred model with 63% vs 24% in Prosit.

To further investigate the difference of CoSpred versus Prosit in full sprectrum and b/y ion prediction, we used an internally generated Hela lysate digestion dataset which was unseen by either model. Both models were applied, and the prediction results of some representative peptides are shown in **Figure 6**. It was expected that fewer peaks would be generated in the Prosit prediction since the model only considers b/y ion fragments. When visualizing all possible fragments such as a/b/c/x/y/z ions, internal fragments, immonium ions, and intact precursor ions, we clearly observed there were many other types of ion fragments besides canonical b/y ions. This result suggests the traditional understanding of peptide spectrum solely based on b/y ion fragmentation may be sub-optimal. To get a higher fidelity spectrum, more ion types should be considered, which is one benefit of utilizing full spectrum prediction framework.

**Figure 5:**
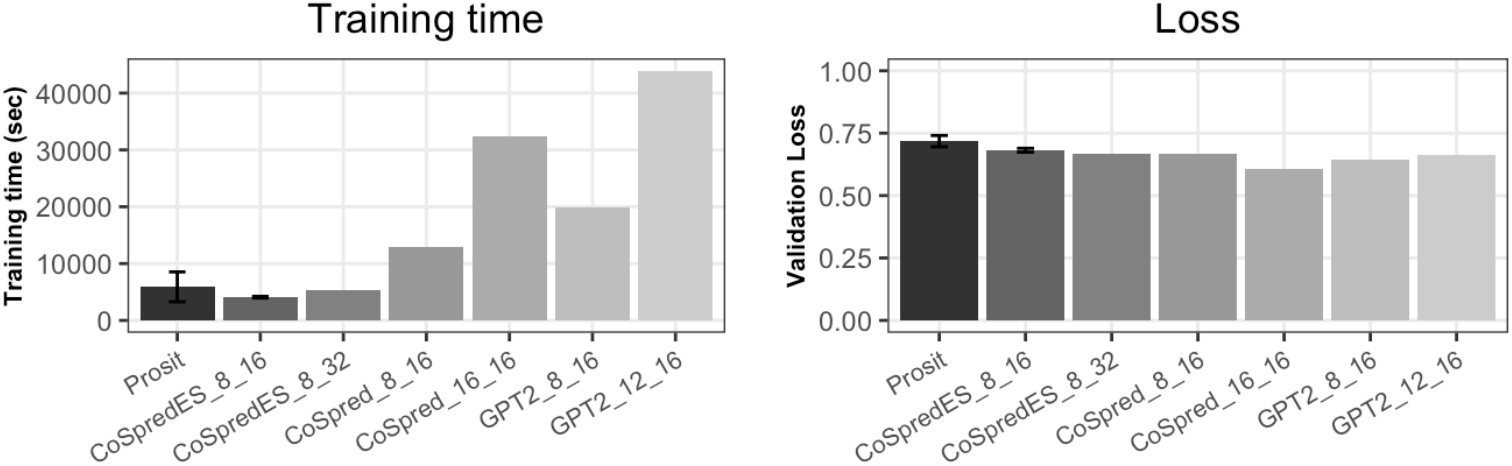
Hyperparameter tuning for full spectrum prediction with the transformer-based architecture. Various layer numbers, head numbers and early stopping were compared in terms of training time in A) and spectra distance loss in B).

**Figure 6:**
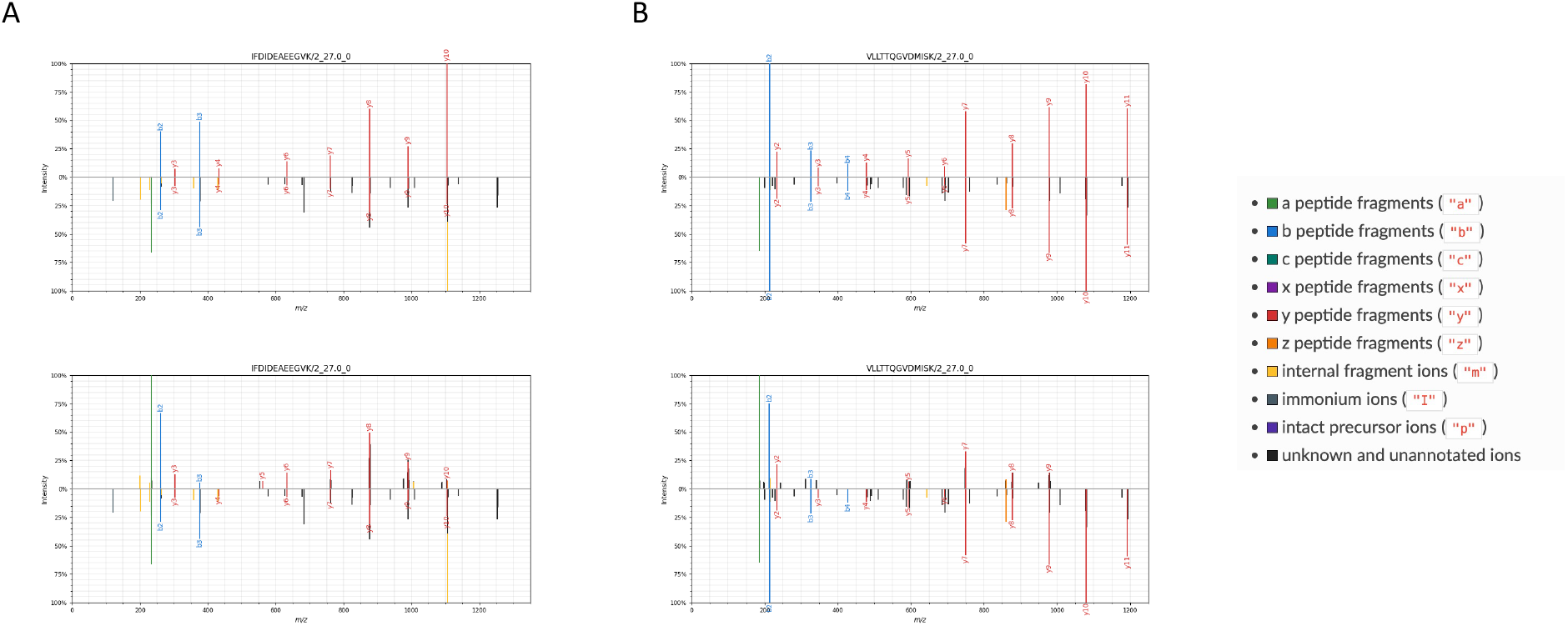
Mirror plots of b/y ion spectrum prediction versus full spectrum prediction of two representative peptides). The upper half of each plot shows predicted spectrum, and the bottom half shows the experimental spectrum. The top two plots were created using the Prosit model, while the bottom two were from the CoSpred model with full spectrum prediction. Column A) and B) represent two different peptides respectively. Note that colors represent different ion types.

### Fine tuning state-of-the-art ML architectures with the CoSpred workflow

Beyond complete spectrum prediction, recurrent neural networks (RNNs) have shown promise in predicting high fidelity spectrum, leading to a revolution of proteomics workflows like data independent acquisition (DIA). However, such ML methods still face challenges with training data size and diversity. For example, the current public Prosit model concentrates on tryptic non-modified peptides with certain types of instrumentation. This may pose problems for highly customized sample preparation and data acquisition methods, where the model performance has yet to be fully evaluated. Here we leverage the flexibility of the CoSpred workflow to retrain the Prosit model using in-house data.

After the Prosit foundation model training process in **Supplemental Figure S6A**, when supplementing an “unseen” dataset to the trained model (about 30,000 PSM), the loss was higher at the beginning (about 0.06 vs 0.02 at the last training epoch of foundation model training). Later, the loss dropped sharply within a few epochs and quickly converged (**Supplemental Figure S6B**), indicating the fine-tuning process was complete for the new dataset in a short period of time.

A representative comparison of the models is shown with various metrices in **Figure 7**. Among all peptides in the test set considered, the overall prediction quality has been improved significantly after fine-tuning, especially for precision of b/y ion prediction (**Figure 7A**) and recall for full spectrum prediction (**Figure 7B**), leading to a higher F1 score. By visualizing some representative spectra, we observed differences in peptide precision and recall before vs post fine tuning (**Supplemental Figure S7A, S7B**). We hypothesize that this improvement for both mass predictions measured by F1 score and intensity prediction measured by cosine similarity is due to the fine-tuning process learning new information about the instrumentation parameters like noise and case specific collision energy, which are not easily apparent from metadata nor have been incorporated in the input features for the foundation model.

**Figure 7:**
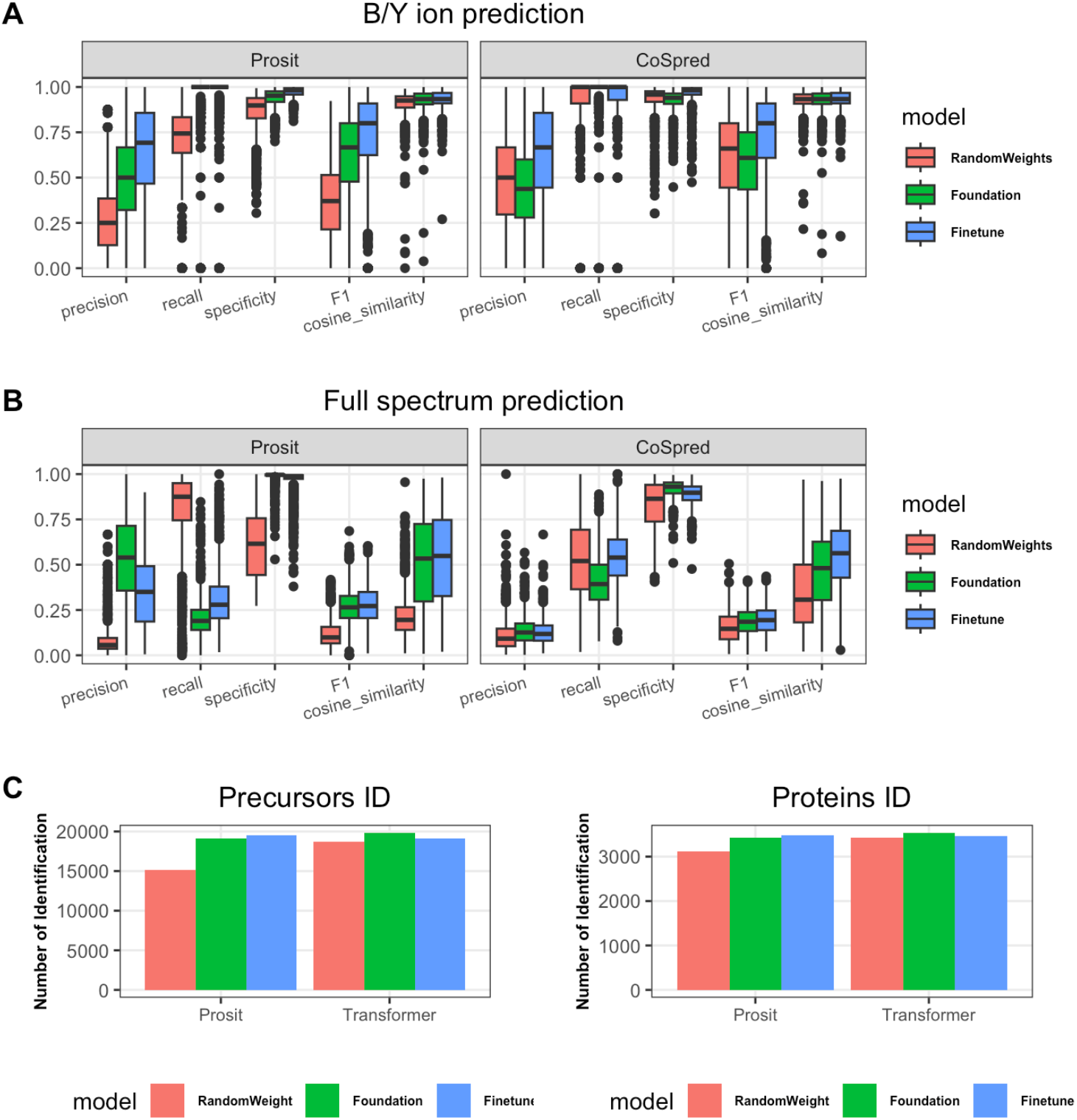
Comparison of fine-tuned vs foundation model for predicted spectrum of all peptides in unseen Hela cell lysate digest dataset. A) and B) are evaluation metrics between b/y ion prediction and full spectrum prediction. C) shows a DIA precursor and protein identification benchmark when applying different training models with Prosit and Transformer architectures.

Interestingly, for training this small set (i.e. train the model with a small number of inputs initiated by random weights), it appears the transformer learns to present meaningful spectrum faster than the Prosit model. The same effect was also observed in fine-tuning scenario, the transformer shows a more significant difference from foundation model, while Prosit model with or without extra tuning gave similar results. On the other hand, both models performed similar for b/y ion prediction regardless of fine-tuning process, indicating the possibility that both deep learning models have extracted most of the information from such task with relatively low complexity.

Furthermore, the 30k PSM used for the transfer learning process, required only about two hours with a regular CPU based laptop (Macbook Pro 2021 model, 8 Core CPU, 64GB RAM) or around 15 mins with A10 GPU, indicating a high feasibility of leveraging this with currently trained large spectrum prediction models and the flexibility of a training workflow which can be optimally applied on user specific prediction tasks.

The most widely used application for such predictions is to create a complete *in-silico* spectral library for data independent acquisition methods (DIA) ^15^ which can be used by software such as DIANN ^15^, MaxDIA ^14^ and Spectronaut^®^ (Bruker Biognosis). Here, we were able to load the fine-tuned spectral library to the DIANN and complete the data processing for internal datasets. Steps for applying models to create spectra libraries for DIA analysis using DIANN can be found in the **Supplementary Materials**. When applying the trained models to a DIA dataset generated from the same instrumentation environment, we observed equivalent or slightly higher identification using the fine-tuned models (**Figure 7C**). Interestingly, the rapid training model (i.e. training a model from random weight initiation with a single raw file) performs better in the transformer architecture than Prosit, which may suggest the possibility to leverage the fast training capability of the transformer for practical cases when a large foundation model is difficult to obtain in the first place.

### Apply fine tuning DL model architectures for post-translational modification proteomic dataset

It was intriguing that we observed promising model improvement using a toy dataset to quickly fine-tune the Prosit model for an in-house dataset which inherits implicit variables like instrumentation and sample preparation protocol, that are not accessible through metadata. We hypothesized it is possible for such fast fine-tuned models to learn novel modifications for peptides that the full model hasn’t been trained on. This would have great potential benefits on applications like biomarker discovery or chemoproteomics where the number of modifications could be high, and the modification type may be unforeseeable. To test this hypothesis, we applied the strategy on in-house generated phosphoproteomics data which has never been seen by the public Prosit model. For performance comparison, a toy model which was initiated with random weights and trained using a single raw file (about 30,000 phospho spectra), as well as the original Prosit model (which was trained with around 7 million non-phospho spectra) were used.

Using the CoSpred workflow with the Prosit model, 1000 spectra were kept as a holdout set. The remainder of the single phosphoproteomics raw file was split into 80% and 20% for training and validation. For the toy model, the training converged after 65 epochs with the loss at about 0.33. In contrast, the original Prosit model had the lowest loss of 0.08. When applying the toy training set to fine-tune the Prosit model, the loss shot up initially, decayed quickly in an early stage, and finally converged at 50 epochs (**Supplemental Figure S8**). The re-training time was about 30 minutes with an A10 GPU.

Comparing the prediction results on the holdout set, we observed consistent patterns for the two models across various peptides. Among a variety of metrices, the fine-tuned model performs best in terms of fragmentation efficiency (precision and recall of correctly predicted peaks) and intensity accuracy (cosine similarity of intensities) (**Figure 8**). Interestingly, the difference between models was more significant in the full prediction workflow. We hypothesize that it may be due to the chemical properties of PTMs influencing non-backbone ions such as neutral loss ions in addition to shifting b/y ions.^38, 39^ Regarding the lower precision measurement, we suspect it was due to small “novel” PTMs which would inevitably contaminate the foundation model which was highly optimized towards un-modified peptide forms. Furthermore, even for modified peptides, a large portion of total fragments are still un-modified fragment ions. The testing result with single phosphorylated peptide (**Supplemental Figure S9A**) versus multiple phosphorylated peptides (**Supplemental Figure S9B**) aligned with our hypothesis. The fine-tuning model has more advantages over the foundation model when the proportion of “unseen” modifications increases.

**Figure 8:**
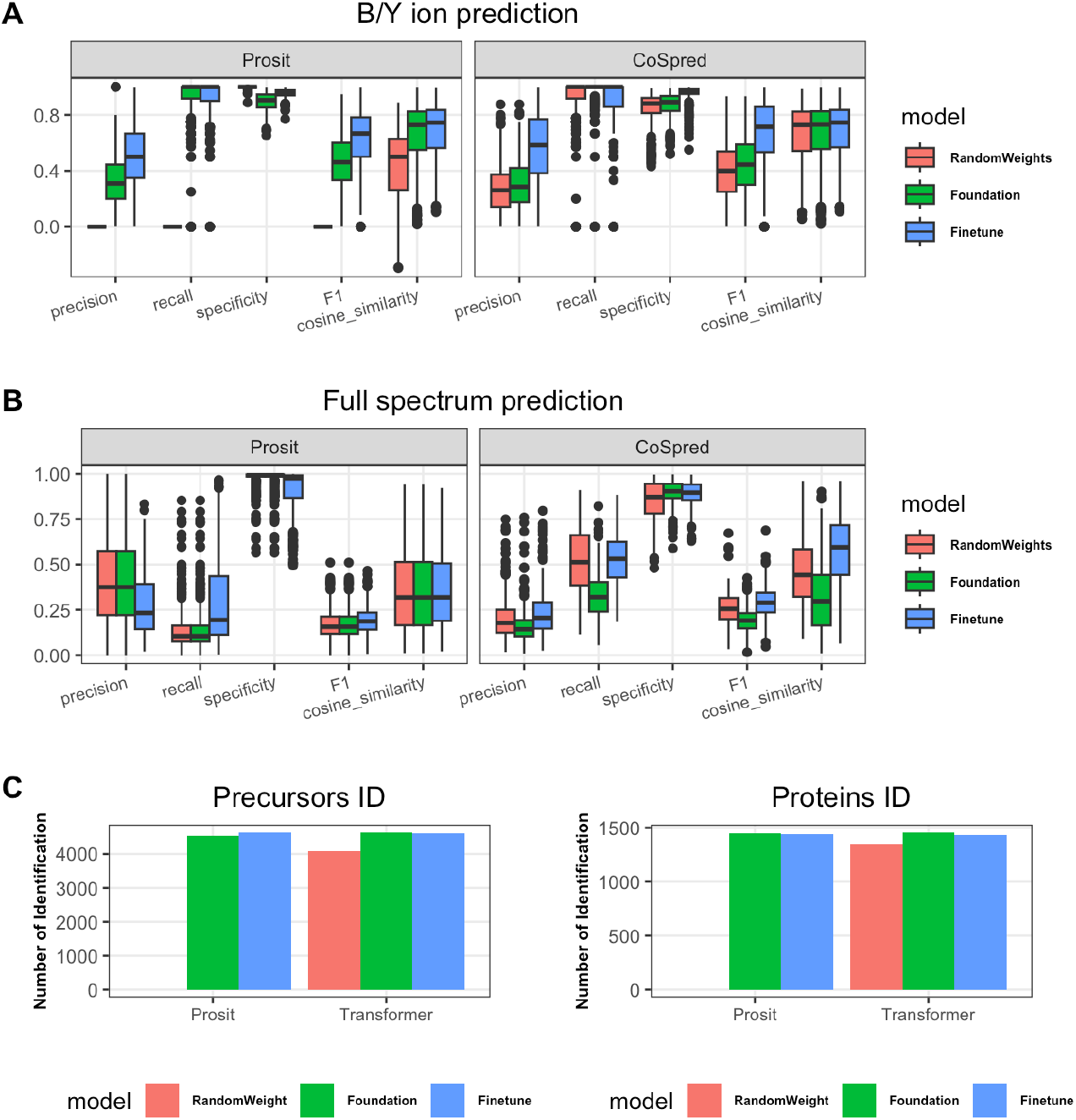
Comparison of fine-tuned vs foundation models for spectra of all peptides in unseen phosphoproteomics dataset. A) and B) are evaluation metrics between b/y ion prediction and full spectrum prediction. C) shows DIA precursor and protein identification benchmark when applying different training models with Prosit and Transformer architectures.

For the benchmarking purposes, we applied the spectra library predicted by the various models to a phosphoproteomic DIA dataset^40^ using DiaNN. Yeast was the species of choice to prevent contamination with spectra in previous training dataset. By comparing the identification of either precursors or proteins, we observed similar performance to the hold-out test set mentioned above, demonstrating transformer-based spectra prediction with fine-tuning capability is an optimal workflow for datasets with novel PTMs.

## DISCUSSION

Backbone ions (a/b/c/x/y/z) have been widely predicted using deep learning algorithms. They typically rely on annotations based on fragmentation rules and use software such as MaxQuant^27^ or Proteome Discoverer. To annotate these backbone ions, the prediction software uses the mass of each amino acid and assigns the fragment ions to these masses. A long-term goal of the proteomics analysis field is the ability to make predictions of spectrum intensities independent of fragmentation rules and to predict complete MSMS spectrum. To date this area has not been fully investigated with only a few studies being published. One of these published studies is the PredFull algorithm, which is based on deep learning methods which use convolutional neural networks to predict both the m/z ratio and its corresponding intensities. The challenging part for this model was creating a framework that worked for all precursor charges. Just to cover charges 1 to 4 it was necessary to add multitask learning on top of the CNN model. In this study, we used a transformer architecture to predict complete spectrum. Using several layers of transformer encoders, the complete spectrum was predicted as a sparse one-dimensional vector of length 3000 with bin width of 0.5Da. This method predicts both m/z ratio and the corresponding intensity and does not require any knowledge about the fragmentation method or fragmentation rules. The input contains the peptide sequence and its precursor charge. CoSpred can predict both backbone and non-backbone ions using this model.

Besides numerical measurement of a prediction, we also applied visualization to evaluate the performance of the transformer-based model. For example, the Spectrum_utils ^31^ package was used to create mirror plots between true and predicted spectrum of a representative peptide is shown in **Figure 6**. The top panel is the predicted spectrum result from the transformer model and the bottom panel is the true spectrum coming from the raw files. Predicted y- and b-ions are shown as red and blue peaks respectively. Other backbone and non-backbone ions are also annotated in different colors. When doing the full spectrum prediction, additional peaks other than b/y ions were also generated and aligned with the reference spectra, suggesting the possibility of using such an architecture for a more comprehensive representation of mass spectrum for a given peptide.

The challenging part of complete spectrum prediction is differentiating valid peaks corresponding to peptide fragments from noise in the spectrum, as valid peaks have a specific pattern, but noise is completely random. To test whether a transformer model that has higher sensitivity at recovering spectrum peaks can still preserve the precision of predicting backbone ions comparable to the other models already available, we compared CoSpred with models predicting backbone ions such as Prosit. That the results were comparable inspired us that a complete spectrum predictor may shed light onto the mechanisms behind the generation of non-backbone ions, an area which is not completely understood. With these preliminary benchmarks, we feel full prediction could be a valuable alternative method for spectrum generation, given its low requirement of prior expert knowledge, faster training procedure, and high sensitivity with balanced performance. In our current implementation, although b/y ion prediction works better for downstream tasks like peptide identification, the data preprocessing procedures like annotating b/y ion is still time consuming. The throughput of the spectrum_util packages for annotating backbone ions is around 10∼100 spectra / second, which quickly becomes computationally expensive for the large datasets easily available for ML training.

In addition to architectures, a user-friendly versatile ML workflow is also in demand. Some published ML models didn’t make the training procedure available to a wide audience, while some of the software had a high dependency on data formats or a strict dependency on the programming environment. Besides investigating novel transformer-based spectrum prediction, CoSpred also aims to provide an end-to-end framework for users to experiment with their own ideas and data in a user-friendly manner. Using this streamlined workflow, we were able to quickly fine-tune and further improve on the well demonstrated Prosit model and get a customized spectra library tailored to internal data types. Furthermore, we also demonstrated the capability of such a workflow towards modified peptides, shedding light on expanding the deep learning-based methods for PTMs and novel modifications when training data is limited.

Future work in complete spectrum prediction is to include other fragmentation methods such as electron transfer/high-energy collision dissociation or ultraviolet photodissociation. With the workflow provided here it will be easy to extend this model for CID or ETD spectra. This workflow can also be extended to non-tryptic peptides commonly coming from immunopeptidomics datasets, which have been shown benefited by high fidelity predicted intensity^35^. Such applications require a large collection of non-tryptic peptides as well as a streamlined training procedure in order to produce an optimized model. Predicted spectrum will also be important for the site localization of PTMs e.g., phosphorylation. DeepPhospho^41^ has created such a deep learning model that integrates spectral library prediction into DIA workflows. However, beyond a very few common modifications like phosphorylation, methionine oxidation, etc, most of modifications occurring naturally or experimentally haven’t been well integrated in the prediction models. We believed our workflow will be a great help for this need. By incorporating preferred specialized models, each one trainable with the optimal training dataset, CoSpred can efficiently accommodate the needs to predict under explored peptides and PTMs.

## ASSOCIATED CONTENT

### Supporting Information

#### Supplemental Figure

Supplemental Figure S1. Charge and length distribution of CoSpred training set.

Supplemental Figure S2. Prediction accuracy with respect to charge and peptide length.

Supplemental Figure S3. Precision recall curve of model training.

Supplemental Figure S4. True vs predicted b/y ion spectrum of two representative spectra.

Supplemental Figure S5. True vs predicted full spectrum of two representative spectra.

Supplemental Figure S6. Loss and performance matrix during the training.

Supplemental Figure S7. Mirror plots of representative peptide spectrum predicted by foundation model and fine-tuned model in Hela lysate digest dataset.

Supplemental Figure S8. Loss dynamics during the training when supplementing the toy training set to trained Prosit model for fine-tune purpose.

Supplemental Figure S9. Mirror plots of representative peptide spectrum predicted by foundation model and fine-tuned model in yeast phosphoproteomics dataset.

#### Supplemental Method

Procedure of using DIANN with predicted spectrum library.

