## Supplemental Methods for "CoSpred: Machine learning workflow to predict tandem mass spectrum in proteomics"

### Procedure of using DIANN with predicted spectrum library

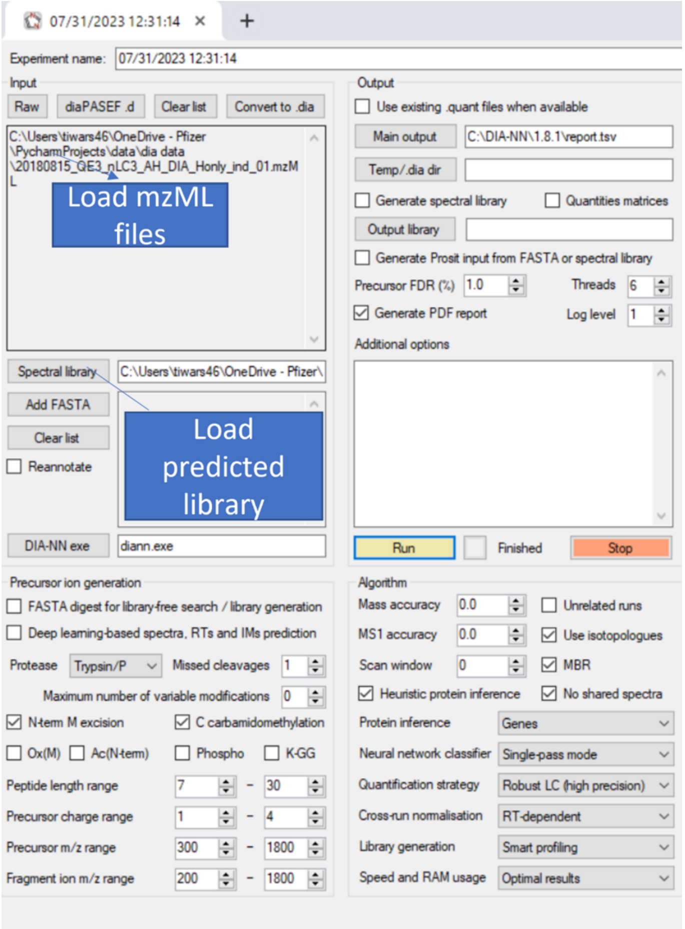


Following these steps to create predicted spectral library.

1. Run training script to train the model
2. Get the best model or saved model in folder cospred models
3. Run cospred_prediction to get the msp format output
4. Use the spectral library in DIANN software
5. In DIANN software, import rawfiles in mzML format, load predicted spectral library and press Run.
