## Supplemental Figures for "CoSpred: Machine learning workflow to predict tandem mass spectrum in proteomics"

### Slide 1
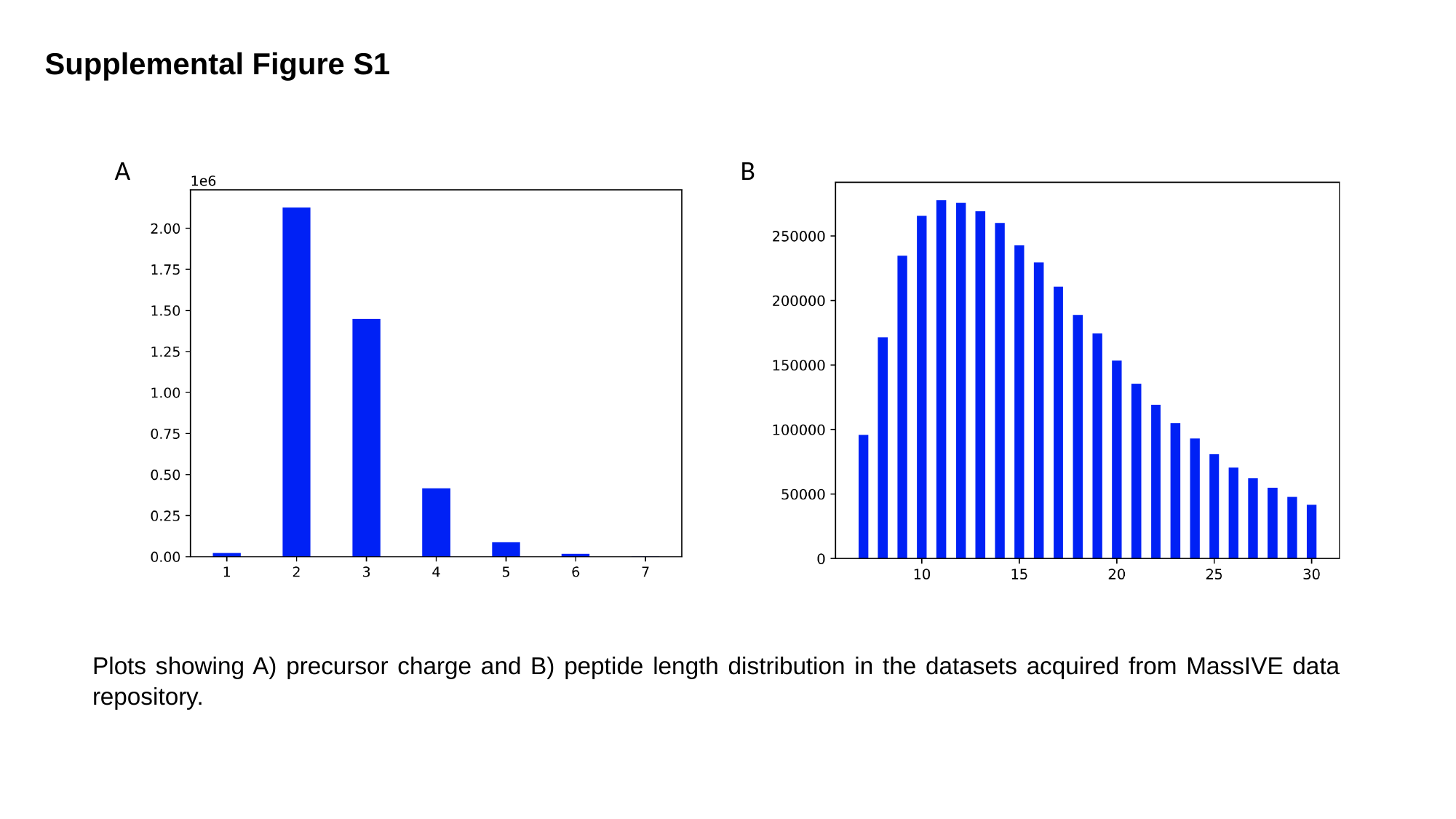

#### Supplemental Figure S1
A
B
Plots showing A) precursor charge and B) peptide length distribution in the datasets acquired from MassIVE data repository.

### Slide 2
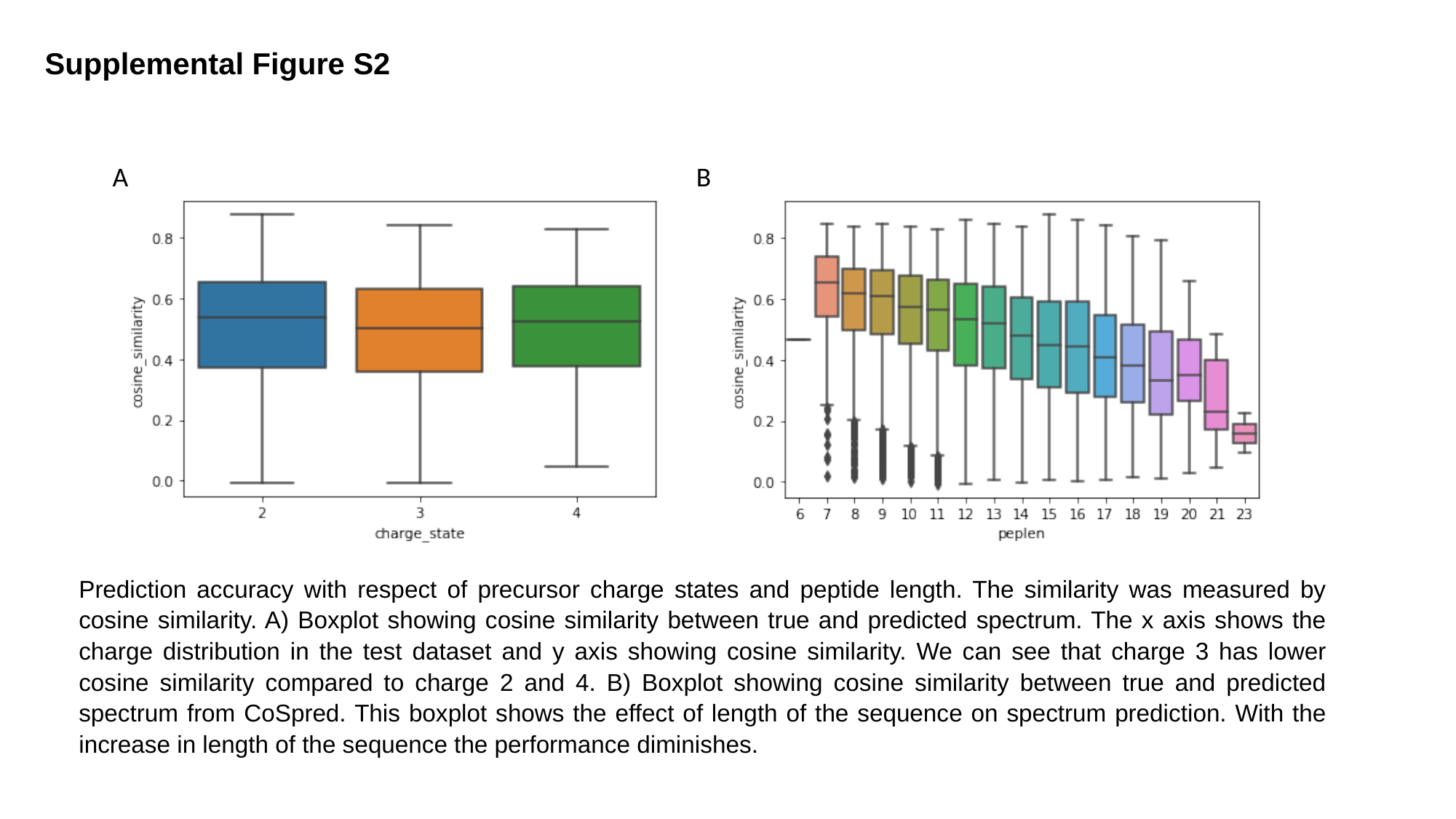

#### Supplemental Figure S2
A
B
Prediction accuracy with respect of precursor charge states and peptide length. The similarity was measured by cosine similarity. A) Boxplot showing cosine similarity between true and predicted spectrum. The x axis shows the charge distribution in the test dataset and y axis showing cosine similarity. We can see that charge 3 has lower cosine similarity compared to charge 2 and 4. B) Boxplot showing cosine similarity between true and predicted spectrum from CoSpred. This boxplot shows the effect of length of the sequence on spectrum prediction. With the increase in length of the sequence the performance diminishes.

### Slide 3
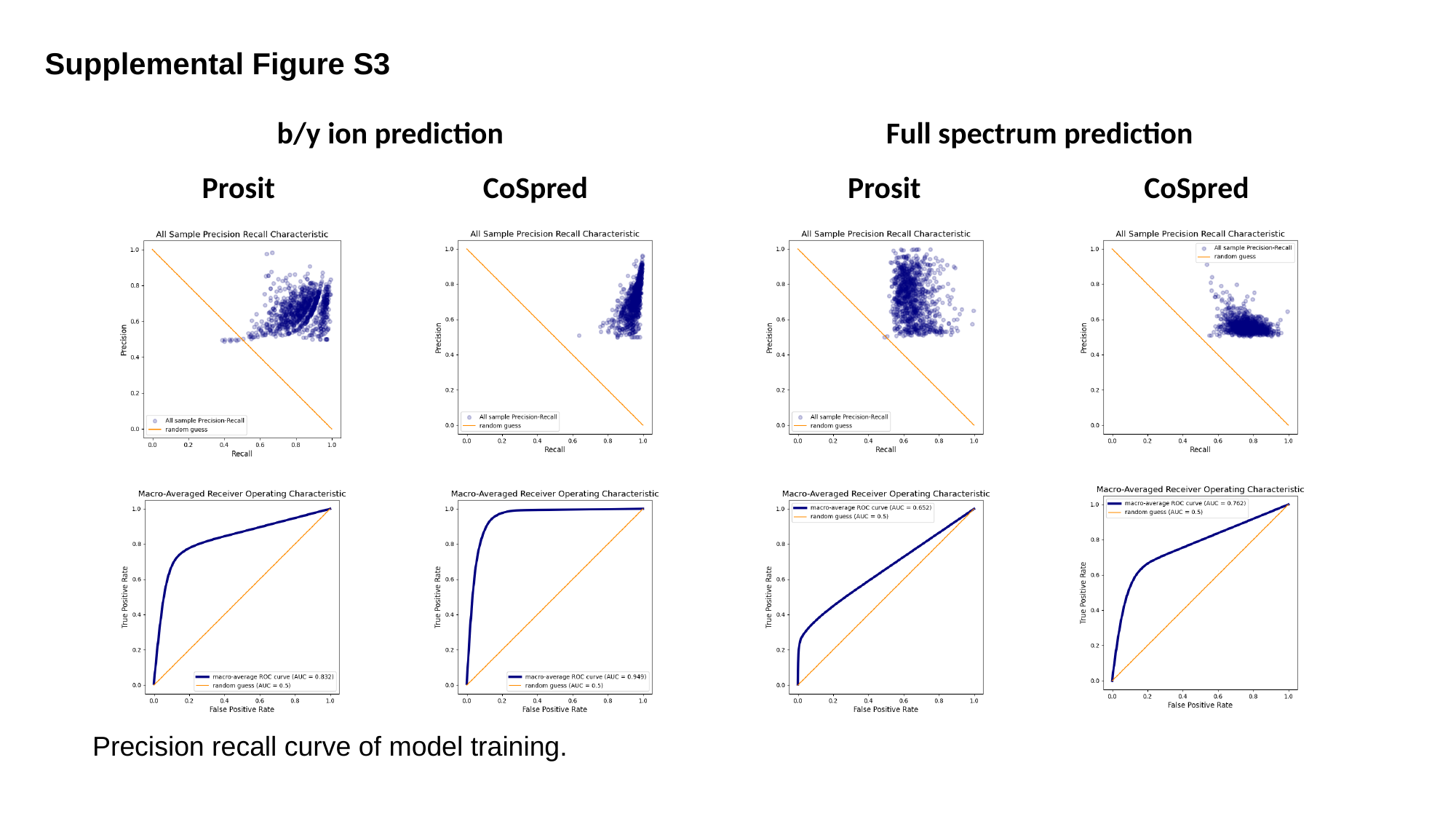

#### Supplemental Figure S3
Full spectrum prediction
b/y ion prediction
Prosit
CoSpred
Prosit
CoSpred
Precision recall curve of model training.

### Slide 4
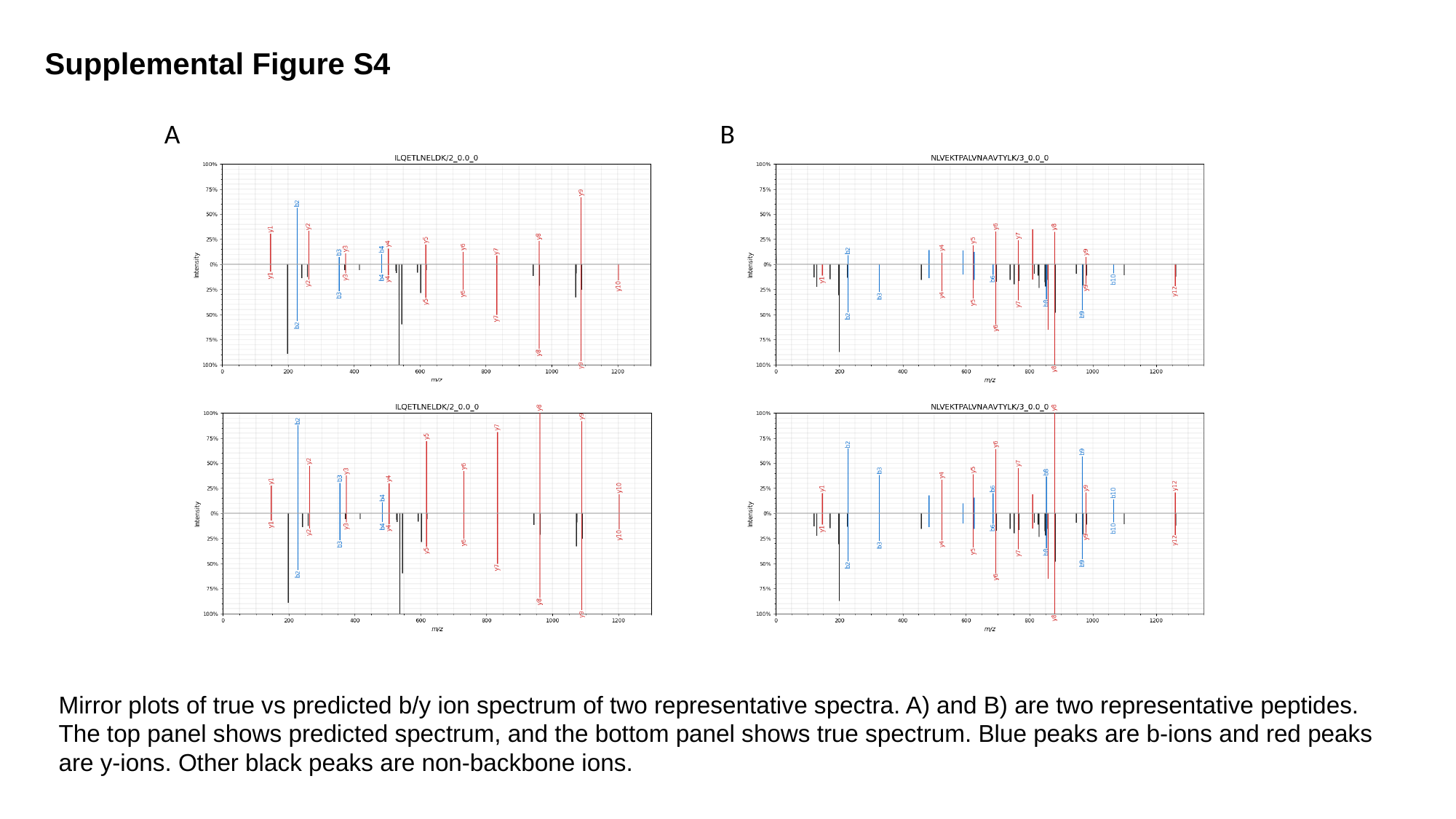

#### Supplemental Figure S4
A
B
Mirror plots of true vs predicted b/y ion spectrum of two representative spectra. A) and B) are two representative peptides. The top panel shows predicted spectrum, and the bottom panel shows true spectrum. Blue peaks are b-ions and red peaks are y-ions. Other black peaks are non-backbone ions.

### Slide 5
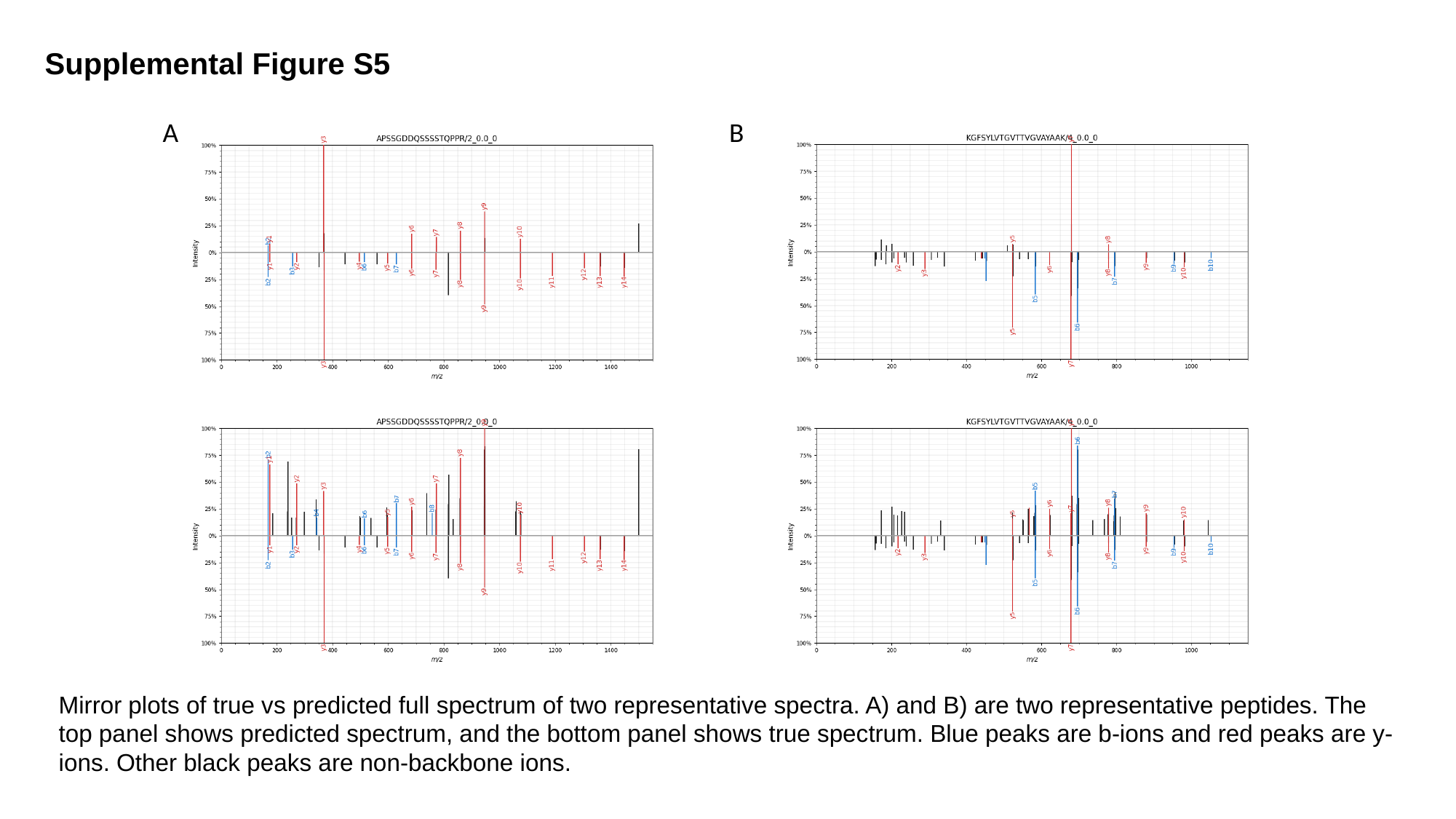

#### Supplemental Figure S5
B
A
Mirror plots of true vs predicted full spectrum of two representative spectra. A) and B) are two representative peptides. The top panel shows predicted spectrum, and the bottom panel shows true spectrum. Blue peaks are b-ions and red peaks are y-ions. Other black peaks are non-backbone ions.

### Slide 6
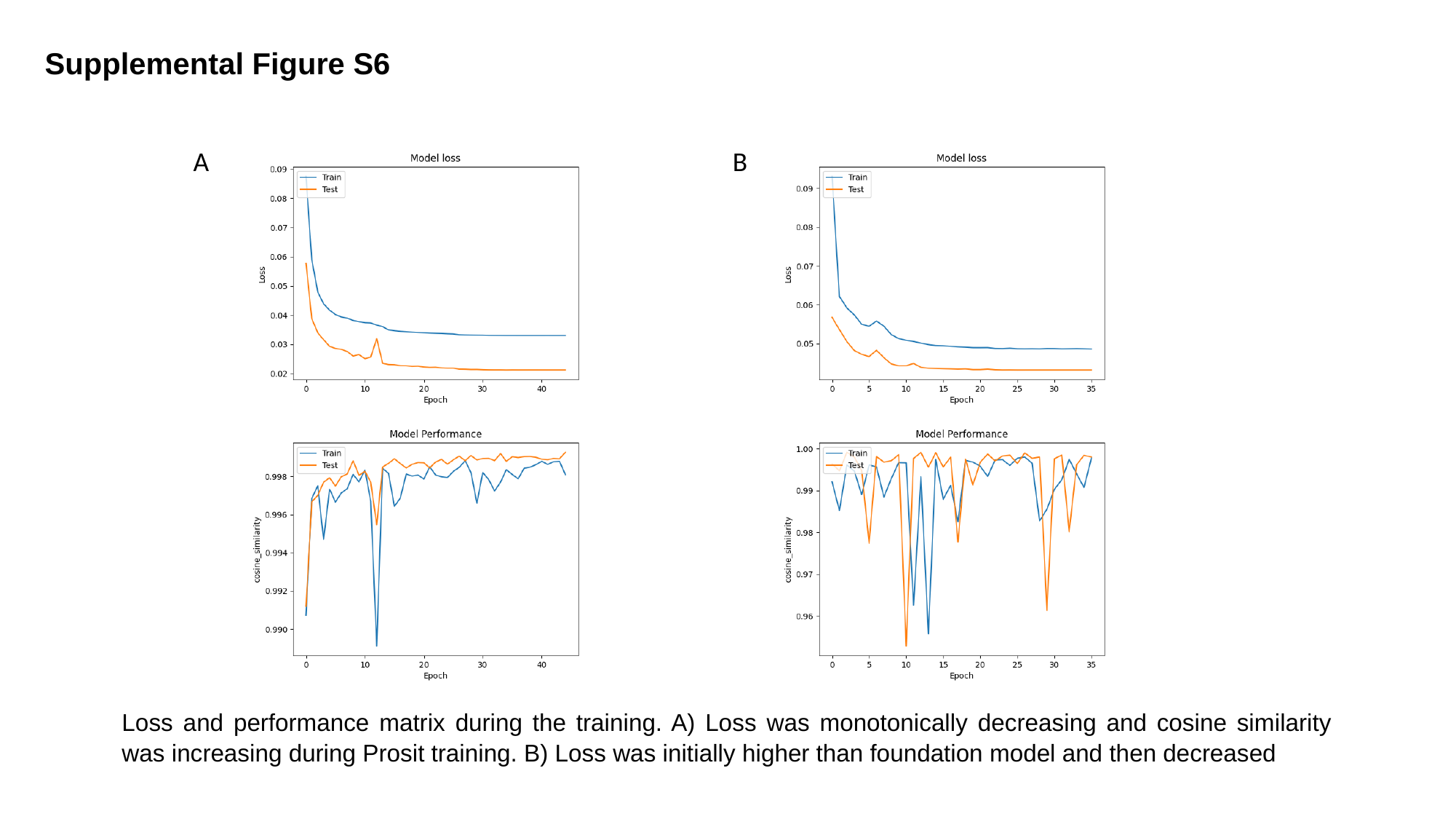

#### Supplemental Figure S6
A
B
Loss and performance matrix during the training. A) Loss was monotonically decreasing and cosine similarity was increasing during Prosit training. B) Loss was initially higher than foundation model and then decreased

### Slide 7
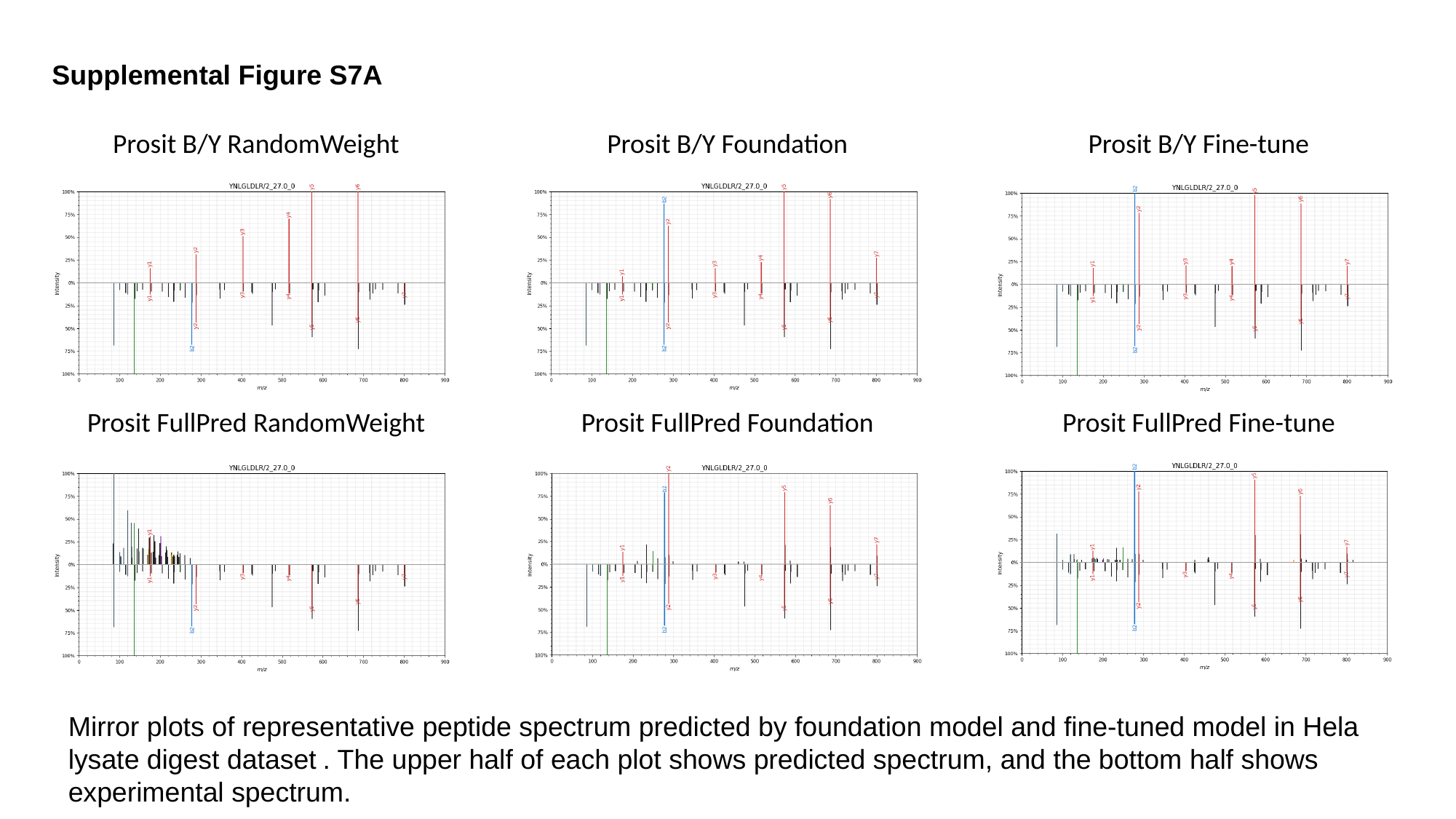

Supplemental Figure S7A
Prosit B/Y RandomWeight
Prosit B/Y Foundation
Prosit B/Y Fine-tune
Prosit FullPred RandomWeight
Prosit FullPred Foundation
Prosit FullPred Fine-tune
Mirror plots of representative peptide spectrum predicted by foundation model and fine-tuned model in Hela lysate digest dataset . The upper half of each plot shows predicted spectrum, and the bottom half shows experimental spectrum.

### Slide 8
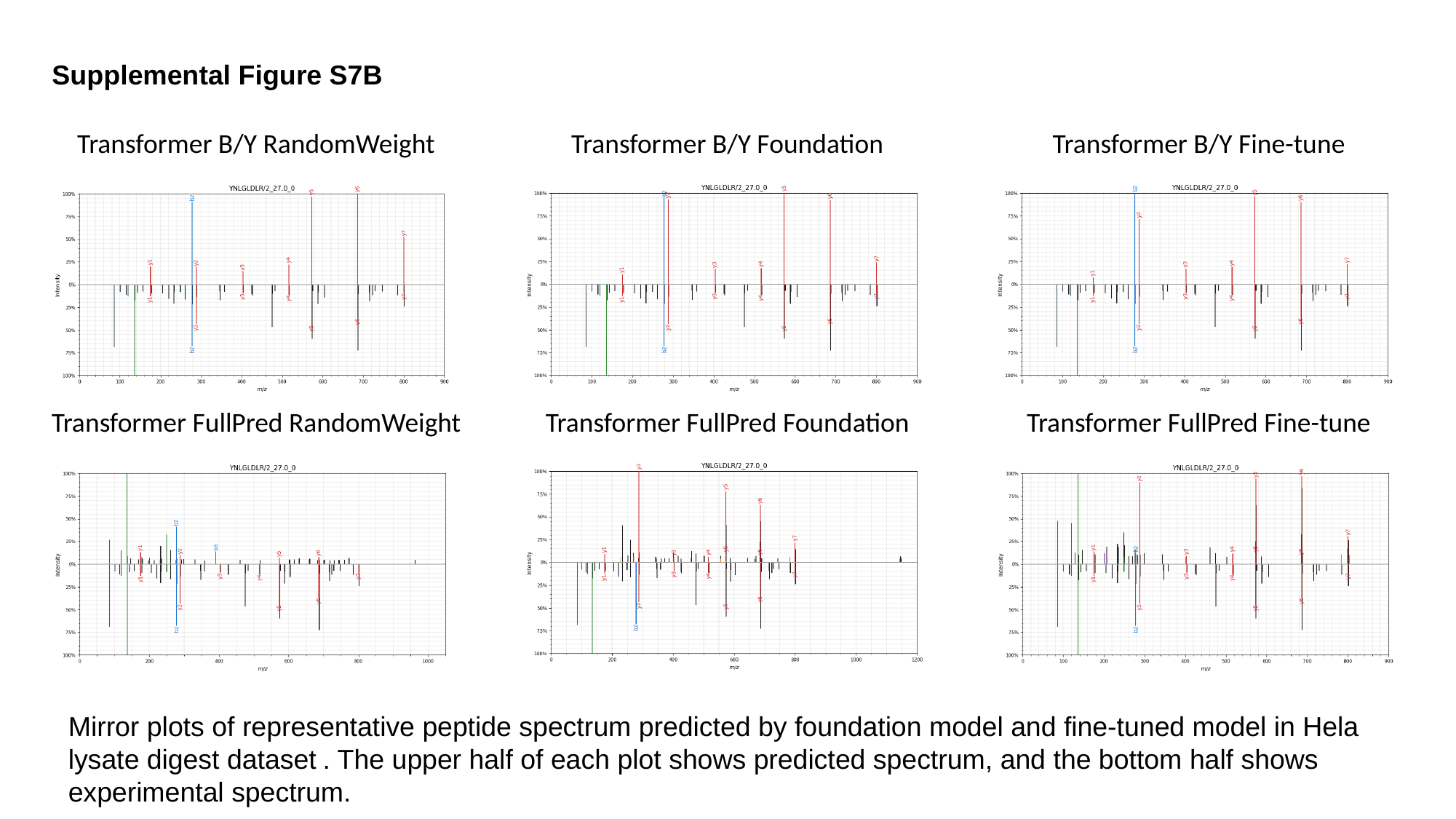

Supplemental Figure S7B
Transformer B/Y RandomWeight
Transformer B/Y Foundation
Transformer B/Y Fine-tune
Transformer FullPred RandomWeight
Transformer FullPred Foundation
Transformer FullPred Fine-tune
Mirror plots of representative peptide spectrum predicted by foundation model and fine-tuned model in Hela lysate digest dataset . The upper half of each plot shows predicted spectrum, and the bottom half shows experimental spectrum.

### Slide 9
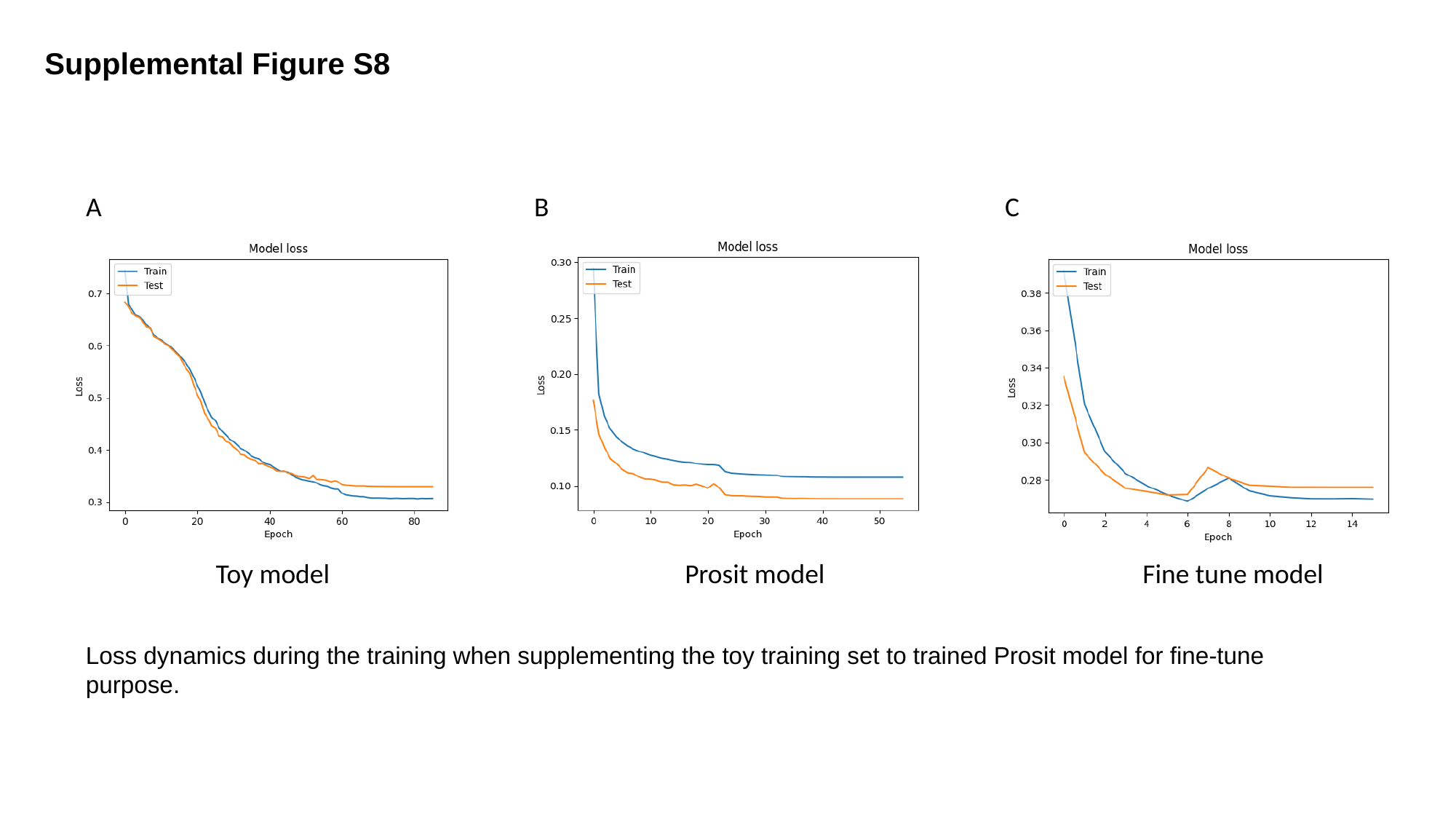

Supplemental Figure S8
A
B
C
Toy model
Prosit model
Fine tune model
Loss dynamics during the training when supplementing the toy training set to trained Prosit model for fine-tune purpose.

### Slide 10
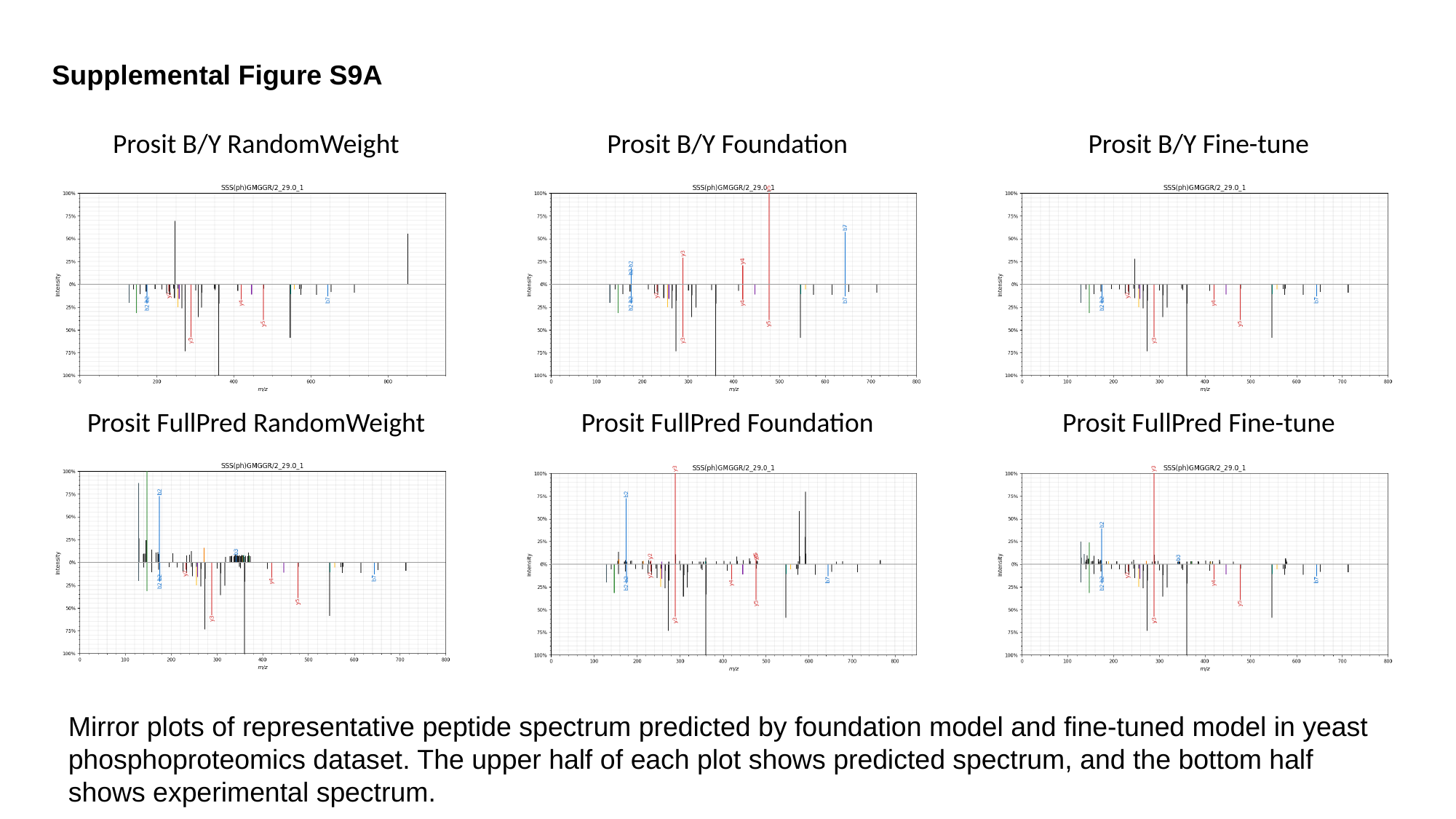

Supplemental Figure S9A
Prosit B/Y RandomWeight
Prosit B/Y Foundation
Prosit B/Y Fine-tune
Prosit FullPred RandomWeight
Prosit FullPred Foundation
Prosit FullPred Fine-tune
Mirror plots of representative peptide spectrum predicted by foundation model and fine-tuned model in yeast phosphoproteomics dataset. The upper half of each plot shows predicted spectrum, and the bottom half shows experimental spectrum.

### Slide 11
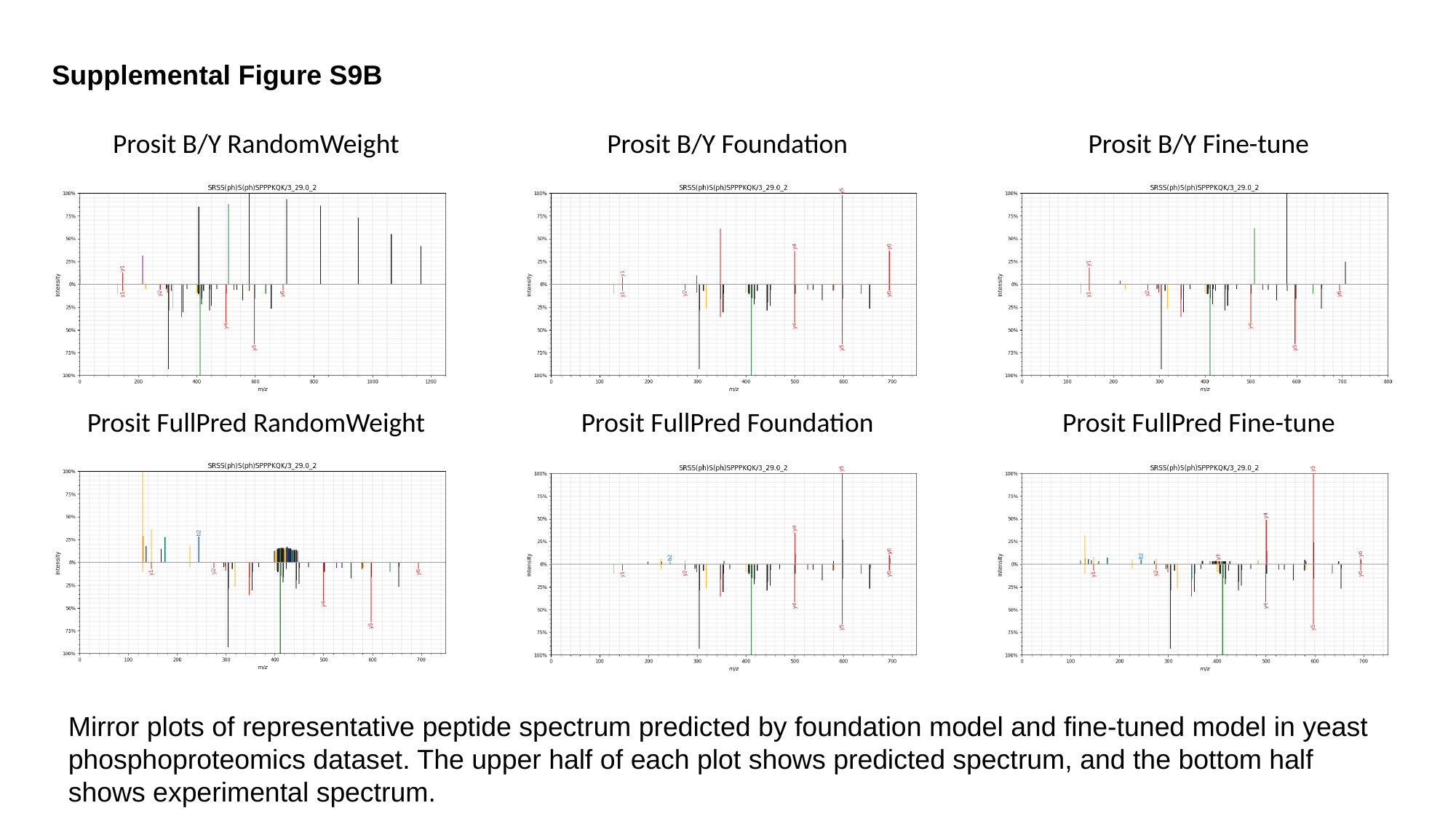

Supplemental Figure S9B
Prosit B/Y RandomWeight
Prosit B/Y Foundation
Prosit B/Y Fine-tune
Prosit FullPred RandomWeight
Prosit FullPred Foundation
Prosit FullPred Fine-tune
Mirror plots of representative peptide spectrum predicted by foundation model and fine-tuned model in yeast phosphoproteomics dataset. The upper half of each plot shows predicted spectrum, and the bottom half shows experimental spectrum.
